# Environmental challenge rewires functional connections among human genes

**DOI:** 10.1101/2023.08.09.552346

**Authors:** Benjamin W. Herken, Garrett T. Wong, Thomas M. Norman, Luke A. Gilbert

## Abstract

A fundamental question in biology is how a limited number of genes combinatorially govern cellular responses to environmental changes. While the prevailing hypothesis is that relationships between genes, processes, and ontologies could be plastic to achieve this adaptability, quantitatively comparing human gene functional connections between specific environmental conditions at scale is very challenging. Therefore, it remains unclear whether and how human genetic interaction networks are rewired in response to changing environmental conditions. Here, we developed a framework for mapping context-specific genetic interactions, enabling us to measure the plasticity of human genetic architecture upon environmental challenge for ∼250,000 interactions, using cell cycle interruption, genotoxic perturbation, and nutrient deprivation as archetypes. We discover large-scale rewiring of human gene relationships across conditions, highlighted by dramatic shifts in the functional connections of epigenetic regulators (TIP60), cell cycle regulators (PP2A), and glycolysis metabolism. Our study demonstrates that upon environmental perturbation, intra-complex genetic rewiring is rare while inter-complex rewiring is common, suggesting a modular and flexible evolutionary genetic strategy that allows a limited number of human genes to enable adaptation to a large number of environmental conditions.

**One Sentence Summary:** Five human genetic interaction maps reveal how the landscape of genes’ functional relationships is rewired as cells experience environmental stress to DNA integrity, cell cycle regulation, and metabolism.

## Main Text

Drawing functional connections between individual genes, pathways, and processes is foundational to our understanding of the interdependence and redundancy of cellular functions. While most studies that seek to nominate putative associations between genes are limited to the basal environmental condition in which their model system is maintained, it is equally important to profile how these associations may change in different cellular states and contexts to adapt the cellular response.

Recently, systems biology approaches have made great progress in characterizing human genes and have provided a wealth of unbiased high-throughput data from which to generate hypotheses (*1–5*). One such approach that provides a scalable snapshot of gene functional relationships is measuring genetic interaction (GI) between gene pairs. Gene pairs that genetically interact are usually found to be related or are components of distinct biological processes whose cooperation is (sometimes unexpectedly) important for cellular homeostasis (*6*, *7*). GI occurs when a combination of genetic perturbations elicits a phenotype that quantitatively differs from an expectation based on the single gene perturbation effects. Classically, GIs are observed in cell viability experiments, where surprisingly deleterious or mitigated growth defects are classified as either synthetic sick/synthetic lethal (SS/SL) or buffering/epistatic interactions, respectively (*8–10*).

GI mapping, in which a large matrix of GIs is systematically and quantitatively measured between sets of genes, enables high-throughput identification of human genetic interactions. In a GI map, each gene’s interaction profile can be used as a multi-dimensional signature to form hierarchical clusters. Clustered genes are often found to exist in protein complexes, have similar roles, or belong to similar ontologies (*11*, *12*). GI mapping has been used successfully in various organisms to assign putative function to uncharacterized genes and discover novel pathways and protein complexes (*5*, *13*). Until recently, technical limitations have precluded large scale measurement of GIs in human cells. The advent of CRISPR-based technology has led to numerous gene deletion, knockdown, and activation screens in human cell culture (*14–19*). These CRISPR methods have been extended to map GIs in human cells and also enable prediction of functional connections through data integration and co-correlation analysis (*20–23*).

While the utility of these analytic frameworks is well appreciated, one persistent shortcoming in their design is the assumption that gene functionality is more or less static and that these approaches implicitly seek a singular annotation to explain gene behavior. Yet, given our current understanding of the size and structure of the human genome, the number of cell types in a human body, and the environmental challenges that must be adapted to at a cellular and organismal level: a one gene - one function relationship is infeasible in humans. The historical difficulty of molecular genetics has necessitated such reductive reasoning, but this could also underlie contradictions in the literature explainable by context specific phenotypes (*24*, *25*).

We posit that the incongruity between the size of the human genome and the myriad conditions it is tasked with responding to necessitate rewiring phenotypes, where a gene product’s functional relationships are environmentally dependent (Figure 1A). No large-scale experiments have investigated the mutability of genetic functional connections systematically in a human cell. Here, for the first time we use CRISPRi-based GI mapping technologies to investigate genetic rewiring in various environmental contexts: S-phase checkpoint inactivation, genotoxic insult from double-strand break formation, and glucose deprivation. We create three environmental and two reference GI maps that exhibit strong, reproducible, and unpredicted phenotypes, revealing gene by gene (GxG) and gene by gene by environment (GxGxE) interactions. Each GI map provides a coherent snapshot of genetic architecture in a particular cell state. We find the composition and intra-cluster interaction profile of established ontologies is broadly conserved across conditions, however the interactions *between* these ontologies are highly context dependent supporting our hypothesis that human genetic interactions are frequently rewired by environmental perturbation. We also observe clusterings of poorly defined genes into new putative complexes, and unpredicted novel GIs between seemingly independent clusters.

**Figure 1.**
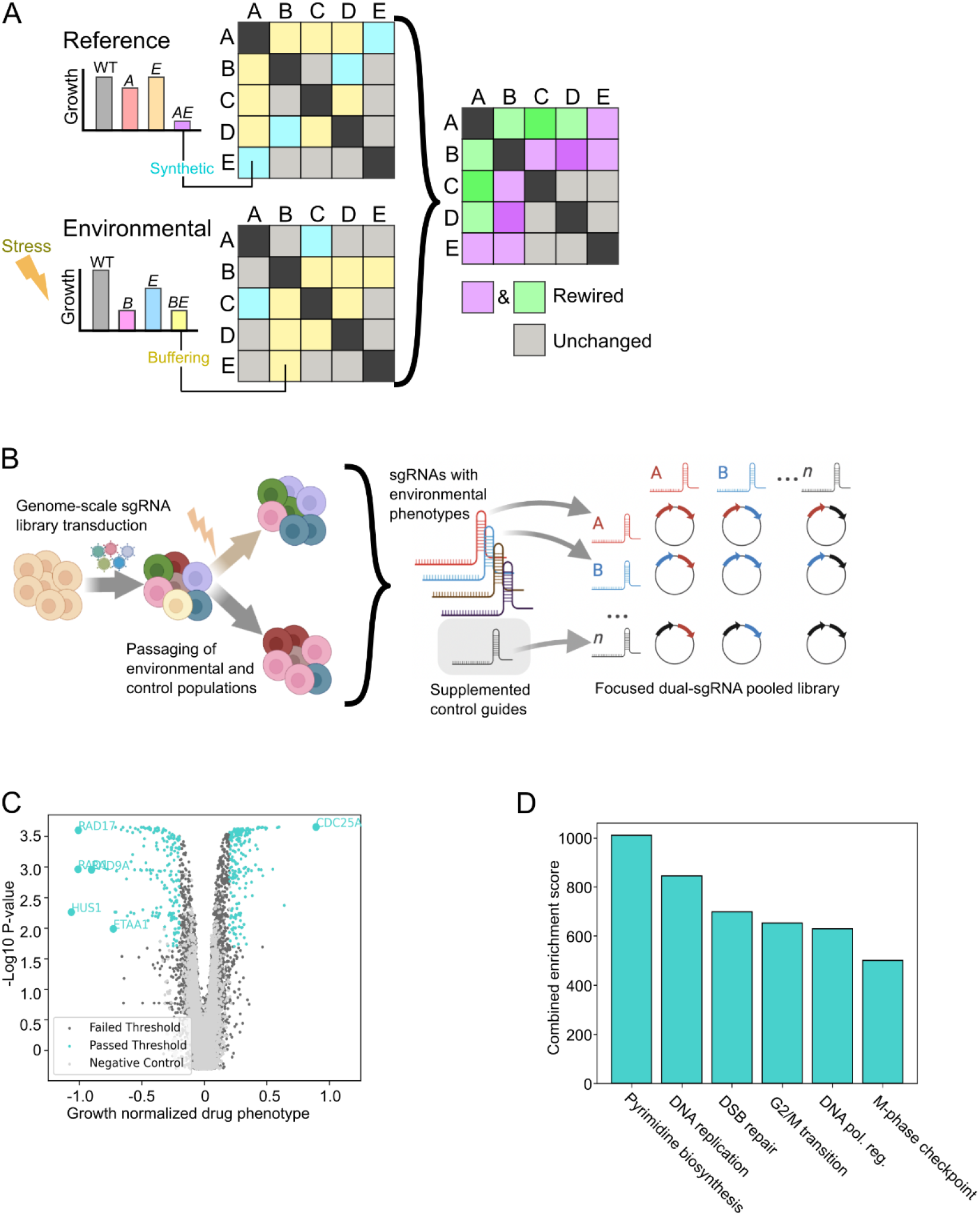
A high-fidelity CRISPRi screen for ATRi modulators. **(A)** Conceptualization of using differential genetic interaction mapping to measure rewiring of gene or ontological functional relationships in response to environmental challenge. Left) Genetic interactions between individual genes under reference and environmental conditions. Cells shaded blue or yellow denote a synthetic sick or buffering interaction, respectively. Middle) A reference GI map (GI) and an environmental GI map (eGI) for a set of genes. Right) A differential GI map (dGI) reveals rewired GIs. **(B)** Experimental workflow for pooled nominating screens leading to a GI mapping library. Cells transduced with a genome-scale lentiviral sgRNA library are passaged with or without environmental perturbation. Conditional phenotypes nominate genes and validated sgRNAs which are then included in a dual sgRNA GI mapping library. **(C)** Volcano plot of gene-level growth normalized drug phenotypes (rho value, see methods) and Mann-Whitney p-value results from nominating genome-scale CRISPRi ATRi screen performed in K562 cells. All 293 threshold passed “hit” genes are highlighted in blue. **(D)** Gene ontology term enrichment from the ATRi nominating screen hit list using the “combined score” metric from Enrichr.

In support of these high level genetic observations, we highlight three biological vignettes wherein the environment dramatically rewires functional connections between biological processes. First, we found that upon S-phase checkpoint inactivation by ATR inhibition, the TIP60 histone acetyltransferase complex radically alters its interaction profile, moving from a peripheral to a key factor in the regulation of core cellular processes such as cell cycle control and metabolism. Second, in response to treatment with etoposide which induces DNA damage, the PP2A complex forms strong pervasive environment specific interactions with a number of ontologies important for DNA replication, fork stabilization, spindle assembly, and translation regulation, suggesting that PP2A may initiate a p53 independent cell death program when replicative damage exceeds the cell’s capacity for repair. From a biomedical perspective, rewired genetic interactions measured between genes implicated in the DNA damage response (DDR) and cell cycle regulation have potential clinical ramifications for patient stratification and the design of new combination therapies, as many of the genes assayed in our experiments have recognized roles in human disease states and both etoposide and the ATR inhibitor we use to induce environmental challenge are approved or in evaluation as anti-cancer therapeutics. Third, we find that upon glucose deprivation the functional connections between glycolytic genes and cellular metabolism are severed, suggesting that enzyme substrate bioavailability strongly modulates metabolic functional connections. In conclusion, our study provides a large dataset of functional connections pertaining to replication fidelity, DNA repair, and metabolism in human cells and illuminates context-dependent rewiring of inter-ontology genetic interactions that reveals new connections between disparate cellular functions.

## Results

### A nominating CRISPRi screen identifies genes that modify sensitivity to ATR inhibition

Two landmark studies have systematically measured environmentally triggered genetic rewiring in yeast, however the nature of this feature of cell biology remains unexplored in human cells (*26*, *27*). In yeast, genetic rewiring phenotypes have been found enriched among genes whose function is related to the cellular response to the environmental challenge in question (*26*). Here, we reasoned that the DNA damage response is a dynamic signaling network which could be enriched for gene functions that can be rewired by environmental perturbations, and therefore decided to focus on this context as a first investigation into rewiring. As a basis to probe rewiring of human gene function, we sought to generate an unbiased list of genes responsible for modifying cellular sensitivity to perturbation of the ATR kinase, a critical regulator of the S-phase DNA damage checkpoint. To this end, we performed a nominating genome-scale CRISPR interference (CRISPRi) screen in K562 human leukemia cells in presence or absence of the ATR inhibitor AZD6738 (ATRi) using our established screening pipeline (Figure 1B and Data S1) (*28*, *29*). Between independent replicates we find highly reproducible sgRNA and gene level phenotypes in both the control and ATRi treated arms of the experiment (Figure S1A&B). A total of 293 genes were found to impart a selective sensitivity or resistance to the ATR inhibitor.

Importantly, many hallmark genes known to be crucial for ATR function or previously found as strong modulators of the ATRi response were identified in our hit gene list (*30–32*). This includes all three members of the 9-1-1 complex (RAD9A, HUS1, RAD1) and its loader RAD17, which appear as the top four genes whose knockdown leads to increased ATRi sensitivity (Figure 1C). We also find CDC25A, whose loss has an established protective effect against ATRi (*30*, *33*), as our strongest resistance hit. The abundance of positive controls among this list as well as high enrichment of gene ontology (GO) terms relevant to DNA repair, cell cycle, and chromatin regulation provide confidence in using this geneset to generate high quality datasets to map genetic interactions (Figure 1D). In addition to predictable DNA repair and cell cycle regulatory factors, our hit list is populated by various genes with GO annotations not canonically associated with ATR biology (Data S1). These surprising additions open the possibility of discovering novel interactions between features of cell biology not usually studied together or mediated by direct physical interactions.

### Reference and ATRi environmental genetic interaction maps of S-phase checkpoint biology

We selected the top two scored sgRNAs targeting each hit gene in our nominating screen to clone a pooled dual-sgRNA GI mapping library (Figure 1B and Data S2), as described previously, containing all combinations of sgRNAs (*20*). Our final library is composed of 408,321 pairs of sgRNAs corresponding to 48,828 unique dual-gene perturbations with eight constructs targeting each gene pair (Figure S1C).

We performed parallel CRISPRi GI screens in K562 cells using our dual-sgRNA library (Figure S1D). One screen was performed under reference (DMSO) conditions, while the other was treated with the ATR inhibitor. Two biological replicates represent each GI experiment (Figure S1E&F).

Our library generates 52 independent measurements of each single gene perturbation phenotype, by pairing every individual gene targeting sgRNA with a panel of non-targeting negative control (ntc) sgRNAs. This enables high quality determination of single gene perturbation phenotypes, which is critical for accurate calling of genetic interactions. The single gene phenotypes derived from both GI screens correlate strongly with those calculated in our nominating screen (Figure S2A-B).

A matrix of sgRNA level genetic interactions was produced using an analytic strategy previously employed for GI mapping in mammalian cells (*20*). For each individual sgRNA, a regression model is applied from the distribution of dual-sgRNA phenotypes that includes the query sgRNA and all single sgRNA phenotypes (Figure 2A&B). GIs of sgRNA pairs that include the query sgRNA are then calculated based on the deviation of the observed pair phenotype from the model. The high representation of ntc sgRNAs paired with each gene targeting sgRNA in our library provides an empirical measurement of noise in our experiments, allowing us to calibrate putative GI scores by normalizing sgRNA level GIs by the standard deviation of the negative control distribution for that specific sgRNA.

**Figure 2.**
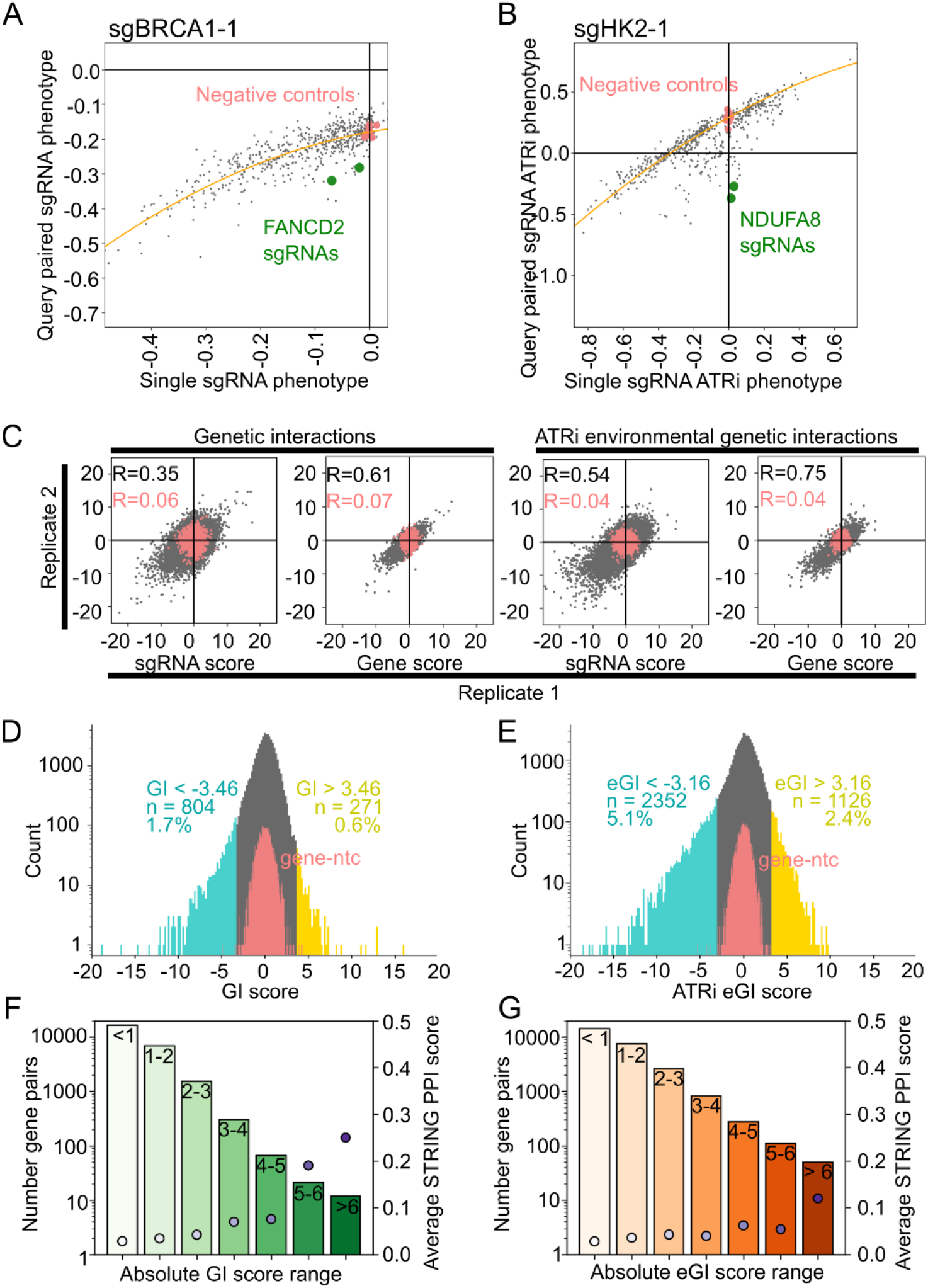
Calling genetic interactions in variable conditions. **(A&B)** Dual-sgRNA phenotype modeling and genetic interaction calling for example query perturbations. BRCA1 reference GI data (A) and HK2 ATRi treated eGI data (B). Negative control distributions for the query sgRNA are highlighted in pink. The model derived from regression of the distributions of single versus query paired sgRNAs is in orange. The two sgRNAs of an example positive control gene anticipated to interact with the query gene are highlighted in green. **(C)** Correlation of sgRNA and gene-level genetic interactions between replicates in the control and ATRi treated conditions. The distributions of non-targeting controls paired with individual sgRNAs or genes are in red. **(D&E)** Distributions of genetic interaction scores called between all gene pairs for reference (D) and ATRi treated (E) conditions. Significance thresholds are represented by histogram shading. Negative and positive GI/eGI scores are highlighted in blue and yellow, denoting synthetic sick and buffering interactions, respectively. Gene-ntc pairs are highlighted in red. **(F&G)** Average STRING protein-protein interaction score for the absolute value reference GI (F) or ATRi eGI (G) scores (dot plot). Population size of each group in each comparison shown as a bar chart.

Finally, gene level GIs are calculated by averaging all sgRNA level GIs that target the same pair of genes. As interactions calculated from the ATRi treated arm of the screen represent a specific gene by gene by environment phenotype, we refer to them as environmental Genetic Interaction (eGI) to distinguish from the DMSO control GI data, which we will hereafter denote as reference GI data (reference GI matrix and ATRi eGI matrix: Data S3&4, respectively).

We find a high degree of correlation between independent experimental replicates for both reference GI and ATRi eGI maps at the sgRNA and the gene level (Figure 2C). We defined the thresholds of significance for GI scores as exceeding four standard deviations of the negative control distribution using the distribution of gene-negative control GIs. This analysis yields 271 (0.6%) positive and 804 (1.7%) negative genetic interactions in the reference condition and 1126 (2.4%) positive and 2352 (5.1%) negative environmental genetic interactions in the ATRi environmental condition (Figure 2D&E). GIs are assumed to be rare and the density of reference GIs found in this experiment is similar to that found in other mammalian GI maps (*20*, *34*). We also find that there is no dependency between each gene perturbation’s primary growth or drug-treated phenotype and the sign or magnitude of its GI or eGI scores (Figure S2C&D), further bolstering confidence in our experimental and analytic approach.

Genetic interactions are often found between genes that physically interact with one another, and we were interested in the extent of overlap between interactions from our dataset and canonical protein-protein interactions (PPI). We used the STRING PPI database to determine how effectively our dataset can recall physical interactions. We find gene pairs that exhibit strong genetic interactions are more likely predicted to physically interact, although the effect is mitigated in the ATRi eGI map and many GIs show no evidence of physical interaction as previously established (Figure 2F&G)(*20*).

Clustering genes by the similarity of their interactions with all other genes in each map provides a scalable and quantitative approach for measuring functional connections, as genes with similar GI profiles tend to be involved in similar biological processes or protein complexes. For each map, we used the average pairwise linkage of the Pearson correlation as a metric to generate hierarchical clusters (Figure 3A&C). This analysis yields robust groupings of genes with known similar functions. Clusters of as few as two genes reveal cognate binding partners (e.g. the relationships between BRCA2/PALB2 and BRCA1/BARD1). Medium sized clusters of between five and twelve genes outline known complexes such as the TIP60 histone acetyltransferase and Complex I of the electron transport chain. Large clusters of up to one hundred genes tend to associate systems level cellular compartments like the mitochondria or concerted cellular bioprocesses like homology directed DNA repair.

**Figure 3.**
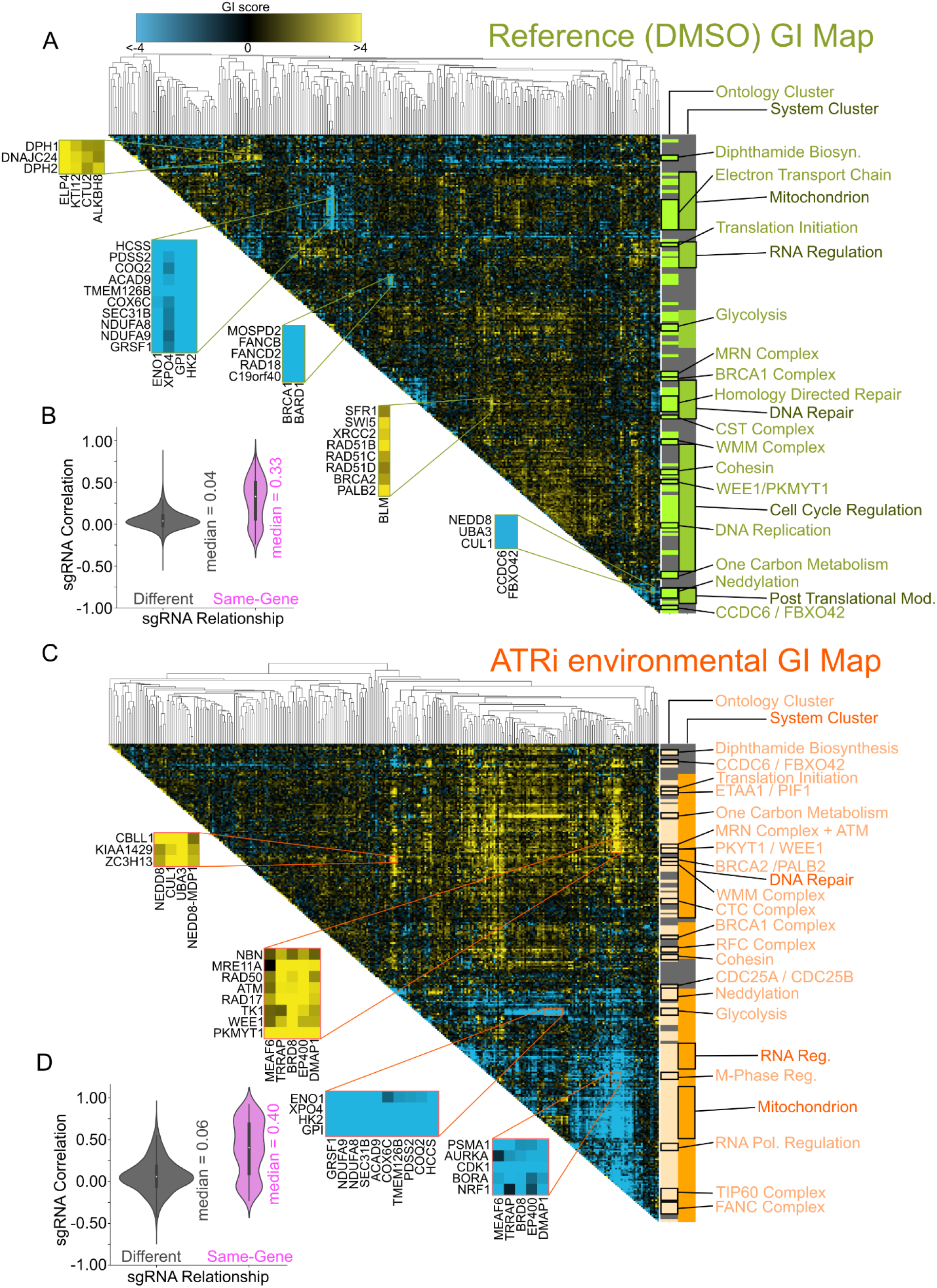
GI and eGI maps of ATR Biology. **(A&C)** Heatmap of genetic interaction scores called for reference GI (A) and ATRi eGI (C) conditions. Cells shaded blue or yellow denote a negative (synthetic sick) or positive (buffering) interaction, respectively (see color bar above 3A). Dendrograms are generated using the average pairwise linkage of the gene level Pearson correlation. Light colored bars to the right of the heatmaps denote clusters that form with high confidence under a stringent threshold (Pearson distance <= 0.55, see methods) that reliably identify discrete ontologies and complexes. Dark colored bars correspond to larger blocks of genes representing cellular compartments or systems-level information that cluster under a relatively more relaxed distance threshold (Pearson distance <= 0.80, see methods). **(B&D)** Violin plots of distributions of Pearson correlations for sgRNA level genetic interactions for different versus same gene targets (gray and purple, respectively) for reference GI (B) and ATRi treated eGI (D) experiments.

In order to determine clustering fidelity we compared the Pearson correlation between the two sgRNAs targeting each gene to the correlation of unrelated sgRNAs. We find in both the reference GI and ATRi eGI maps that same-gene sgRNAs correlate well (Pearson R = 0.33 and 0.40 for GI and eGI sgRNA level maps, respectively), whereas sgRNAs targeting unrelated genes are uncorrelated (Pearson R = 0.04 and 0.06 for GI and eGI sgRNA level maps, respectively) (Figure 3B&D).

Providing further confidence in our analytic framework, a subset of GIs measured in both environmental conditions were validated by fluorescence competition assays. In these assays, perturbed and wild-type cells are grown together and fitness phenotypes are calculated by monitoring depletion or enrichment of the perturbed cells over time using fluorescent protein expression as a proxy to mark genetically perturbed cells (Figure S3A-D). Additionally, GIs and eGIs measured in K562 cells were conserved in A549 lung cancer cells by fluorescence competition assay, suggesting that at least within DNA replication and S-phase checkpoint biology, cell line specific interactions are likely uncommon (Figure S3E-H)

Importantly, the GIs we observe recapitulate established functional relationships, such as the interaction between the BRCA1 and Fanconi Anemia (FA) complexes (Figure 3A)(*35*). Intriguingly, this interaction is not conserved in the ATRi eGI map, suggesting context specific GIs are measurable using this analytic framework and that loss of ATR function might sever the connection between sensing of a lesion and recruitment of effector proteins to manage its repair.

Measuring genetic interactions involving essential genes has historically required complicating hybrid approaches to perturbing gene function. However, knockdown by CRISPRi enables high quality determination of GI with genes whose function is critical to cellular homeostasis. For instance, robust GI and eGI profiles were measured for PPP2R1A, an essential gene and component of the Protein Phosphatase 2A (PP2A) complex that restricts cell cycle progression through multifaceted mechanisms. We detected negative genetic interactions in the reference GI map between PPP2R1A and WEE1, PKMYT1, and MDC1 - all important checks on ectopic progression into M-phase (Data S3)(*36*, *37*). These negative GIs may be indicative of a compounding failure to regulate cell cycle progression leading to mitotic catastrophe.

Predictable clusters are helpful for orienting our understanding of the maps into echelons of biological complexity but are perhaps most interesting when they include understudied genes that cluster with one another or with established complexes, as this allows rapid generation of hypotheses regarding the functions of these genes. Interestingly, we observed that the karyopherin XPO4 clusters closely with the dedicated glycolysis factors GPI, HK2 and ENO1 and all four genes display strong negative GIs with mitochondrial factors involved in oxidative phosphorylation in both reference and ATRi eGI maps (Figure 3A&C). These interactions were validated by a fluorescence competition assay (Figure S3C&D), suggesting a specific role for nuclear-cytoplasmic translocation in the function of glucose metabolism, or a hitherto unidentified function for XPO4.

Another example of novel clustering is illustrated in a grouping containing only CCDC6 and FBXO42. CCDC6, a gene that is mostly known for its propensity to fuse with proto-oncogenes in various human malignancies, is also thought to have a role in potentiation of DNA damage signaling through the phosphorylated histone variant γH2AX, though a direct mechanism for this activity remains unclear (*38*). FBXO42 belongs to a family of genes that provide target specificity to SCF ubiquitin ligase complexes. This cluster negatively interacts in the GI and eGI maps with SCF core and accessory proteins such as CUL1, UBA3, and NEDD8 (Figure 3A). Both genes also display severe negative GIs with PPP2R1A, suggesting that CCDC6 and FBXO42 may be influencing cell cycle progression via a mechanism involving PP2A biology (Data S3). Supporting a function for these genes as cell cycle regulators, CCDC6 or FBXO42 knockdown in ATRi-treated K562 cells resulted in a significant depletion of cells in G1 phase, as measured by propidium iodine staining, while no such cell cycle disruption was observed in control cells (Figure S2E-G). Taken together these lines of evidence suggest a model in which CCDC6 and FBXO42 might serve as a check on progression from G1 phase likely mediated by SCF dependent degradation of target proteins. The perturbation of these genes may trigger further ectopic cell cycling that negatively synergizes with ATR inhibition to dramatically accelerate the cell cycle, leading to mitotic catastrophe..

### Differential interactions reveal rewiring of inter-complex functional connections

Inspection of our data revealed a large degree of contrast in GIs between the reference and ATRi maps, which surprised us (Figure 3A&C). We first sought to understand if the large number of environmental GIs was due to accentuation of existing phenotypes from the reference map, or new genetic interaction due to broad rewiring of genetic relationships in the treated condition (Figure 4A). The GI and eGI map scores correlate (Pearson R = 0.42), but sizable populations of interacting gene pairs are only found in one condition. Many interactions measured in the eGI map are not detectable in the GI map, even when accounting for variations in signal strength between conditions (Figure 4B). These observations led us to posit that we can quantify differential GIs (dGIs) between environmental conditions and that these differential interactions might represent genetic rewiring, which has not been systematically studied in human cells.

**Figure 4.**
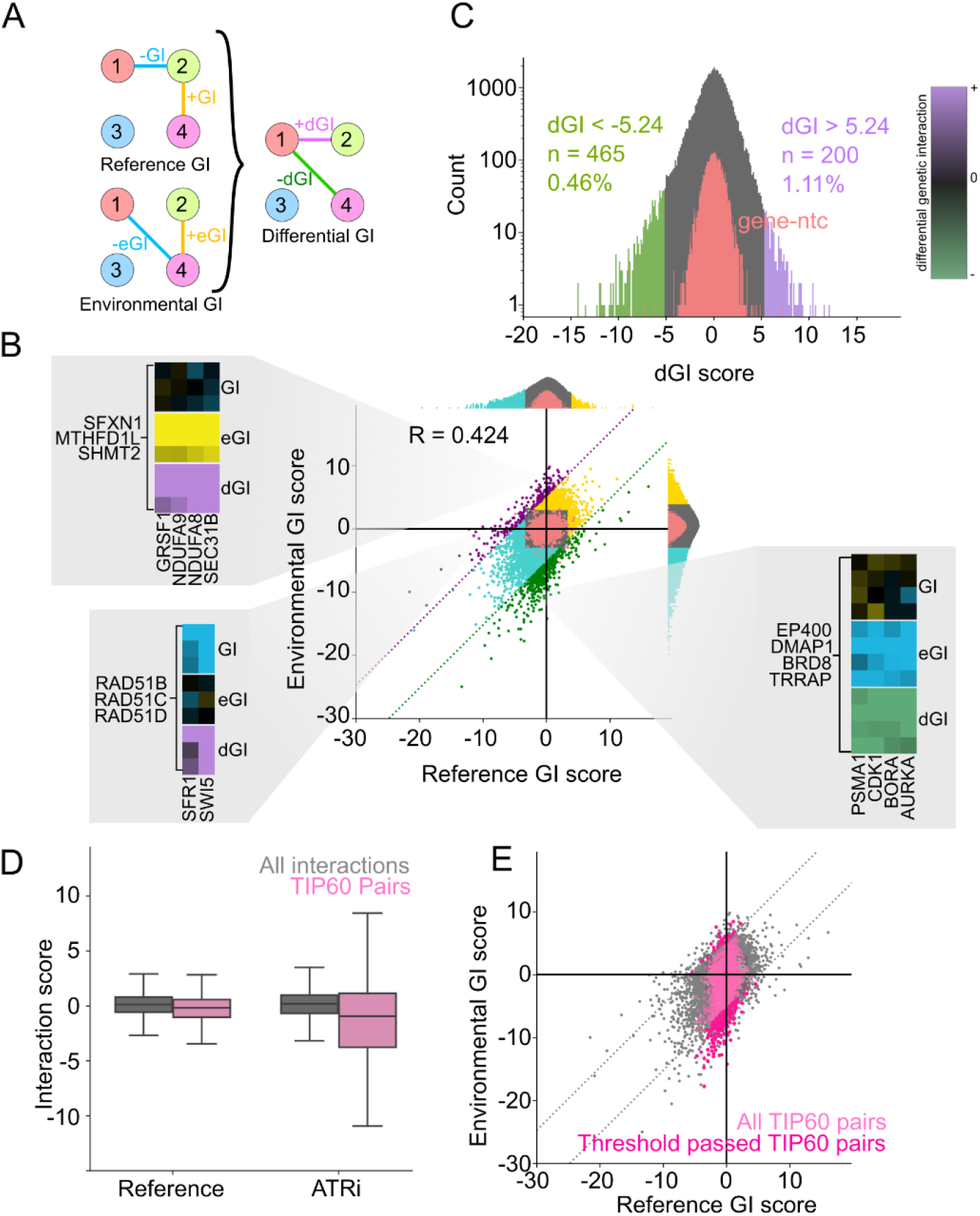
Analysis of ATRi differential genetic interactions. **(A)** Diagram for calculating differential genetic interactions (dGI) from environmental genetic interactions (eGI) and reference genetic interactions (GI). **(B)** Scatter plot of all gene level reference GIs and ATRi eGIs. Significance thresholds are denoted by dotted vertical, horizontal, and diagonal lines for GI, eGI, and dGIs, respectively. Blue or yellow regions of the plot or cells in cutouts denote synthetic sick or buffering interactions, respectively. **(C)** Distribution of dGI scores with significance thresholds denoted by histogram shading. Negative and positive threshold-passed dGI interacting gene pairs highlighted in green and purple, respectively. Gene-ntc pairs are highlighted in red. **(D)** Boxplot of reference GIs and ATRi eGIs for TIP60 genes (pink) relative to all interactions (gray). **(E)** Scatterplot from Figure 4C but with all measured TIP60 gene interactions highlighted in pink. Threshold passed dGIs are shown in darker pink.

To begin to examine the features of this potential biology, we created a third matrix by taking the difference of each gene pair’s score in the ATRi eGI and reference GI maps (Data S5). In this matrix, interactions that are refractory to ATRi are removed, and the resulting matrix displays only the extent to which genetic interactions are altered by treatment with ATRi (Figure 4A). We defined the thresholds of significance for dGI scores as exceeding five times the standard deviation of the distribution of gene-ntc dGI scores. Additionally, we do not consider threshold passed dGIs resulting from eGI/GI scores where neither constituent score is considered significant. We find 200 positive and 465 negative differential genetic interactions with this methodology (Figure 4C).

Analysis of dGIs provides a means of measuring conditionality in relationships of genes and ontologies and draws attention to interactions that, in absence of the comparison, might not invite scrutiny. For example, in the reference GI map, the RAD51 paralogs (RAD51B, RAD51C, RAD51D) cluster with other genes with well established ties to homology directed repair including SFR1, and SWI5 (Figure 4B). This group displays negative GIs amongst themselves, illustrating an important interdependence in responding to this highly cytotoxic type of DNA lesion. In the ATRi treated environmental map, clustering of the complex is maintained but its relationship with other aspects of DSB repair is interrupted. Many DSB repair sub-complexes lose their inter-cluster interaction profile and generally display many fewer eGIs with other ontologies. This trend, in which an interaction found in a control condition is lost upon treatment, sometimes called a “masked” interaction (*26*), suggests an inability to mobilize effective HDR when ATR is inhibited, consistent with our current understanding of the DDR.

In another example of rewiring, we find a conserved cluster of three genes (MTHFD1L, SFXN1, and SHMT2), implicated in one carbon metabolism, that conditionally have a strong positive genetic interactions with most genes in the mitochondrial supercluster in the ATRi condition but not in the reference condition (Figure 4B). These genes are notable in that they all are specifically implicated in a step of folate metabolism localized to the mitochondria. The conditional nature of this interaction, and the shared subcellular localization of proteins derived from both these clusters, suggests that destabilization of mitochondrial homeostasis and energy production is counter-intuitively an important avenue to resist the aberrant cell cycle effects of ATRi. Perhaps slowing cell growth and division through energy starvation and allowing more time to sufficiently replicate and repair DNA, acts to limit erroneous progression into M-phase, thus preventing mitotic catastrophe.

### The TIP60 HAT complex displays extensively rewired GIs

One trend we observed in our data is that many gene pairs with negative eGIs in the ATRi treated map - indicating synthetic lethality - do not interact under reference conditions. On closer analysis, a striking number of these pairs contain at least one component of the TIP60/NuA4 HAT complex (Figure 4B,D&E). TIP60 is a highly conserved family of genes with diverse roles primarily in stimulating gene expression, mobilizing the DDR through ATM activation, and facilitating histone variant exchange, particularly H2AZ and H2AX (*39–41*).

Many, but not all, annotated members of TIP60 are present in our dataset (DMAP1, EP400, BRD8, MRGBP, EPC2, and TRRAP). All six of these genes cluster in the GI map however only a conserved core of four (BRD8, DMAP1, EP400, and TRRAP) cluster in the ATRi eGI map. This minor variation in clustering may support the notion of sub-complex specialization that has been reported in the literature (*42*).

A closer look at the TIP60 perturbation growth phenotypes in the ATRi condition shows that TIP60 knockdown by itself provides a protective effect against ATRi, as was observed in the nominating screen. However, when TIP60 knockdown is paired with many other sgRNAs, the dual sgRNA perturbation results in markedly sicker cells. Our data implicate TIP60 as important to the cellular response to replication stress in two ways. First, loss of TIP60 confers a selective advantage upon inhibition of ATR. Second, S-phase checkpoint inactivation results in a cell state in which TIP60 forms genetic interactions with many key gene sets that themselves present an ATRi specific phenotype upon perturbation. These interactions tend to be associated with core cellular processes, like regulators of metabolism, gene expression, and cell cycle progression rather than DNA damage specific effectors.

We next investigated if the strong growth and interaction phenotypes observed from loss of TIP60 in the context of ATR inhibition caused similarly unexpected alterations in gene expression, reasoning that TIP60’s annotated role as an important regulator of transcription might make it particularly amenable to this type of phenotypic readout. To this end we employed RNA sequencing (RNAseq) to measure genes that are differentially expressed upon knockdown of TIP60. We find that knockdown of either the BRD8 or DMAP1 subunits of TIP60 induce widespread changes in gene expression relative to unperturbed K562 cells in the control condition, underscoring the importance of this complex. As expected, we find considerable overlap in differentially expressed genes following knockdown of either BRD8 or DMAP1 (Figure 5A).

**Figure 5.**
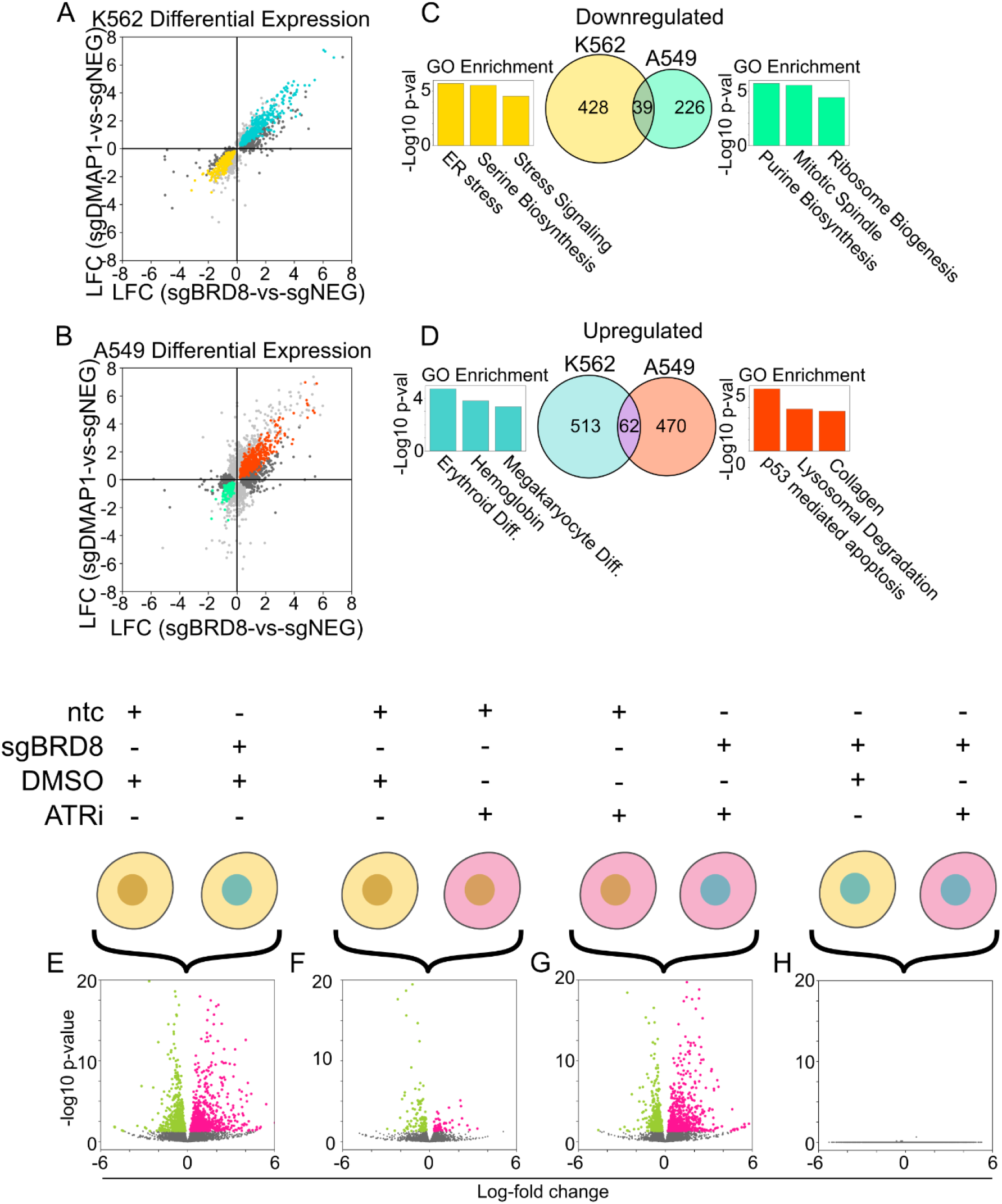
Analysis of the effects of TIP60 knockdown on gene expression and DDR activity. **(A&B)** Scatterplot of log fold change (LFC) in gene expression in K562 or A549 cells upon perturbation of either BRD8 or DMAP1 relative to non-targeting control (ntc). Genes in dark gray and light gray are found to be differentially expressed (adjusted p-value < 0.05) specifically in the BRD8 or DMAP1 perturbations, respectively. Colored genes are significantly differentially expressed in both perturbations. **(C&D)** Venn diagrams of overlap in differentially expressed genes that are regulated by both BRD8 and DMAP1 perturbation between cell types. Gene ontology term enrichment derived from each grouping of genes by Enrichr. **(E-H)** Volcano plots of four differential expression comparisons in K562 cells upon perturbation of BRD8, treatment with ATRi or both.

We repeated this experiment in A549 cells to understand if we could extrapolate these findings in another cell line and observed a similarly large number of differentially expressed genes upon TIP60 perturbation (BRD8 or DMAP1 knockdown) (Figure 5B). Although differentially expressed genes in the two TIP60 perturbations are less correlated in A549 than the same comparison in K562, this correlation is stronger than that of perturbations between cell lines (Figure S4A-H). Accordingly, the regulation of specific genes between the K562 and A549 arms of the experiment was considerably different. GO term enrichment of genes differentially expressed exclusively in A549 show clear activation of a p53 dependent apoptotic program, with a simultaneous downregulation of key markers of cell growth and homeostasis, including mitotic factors and ribosome biogenesis. In contrast, in K562, which does not have active p53 signaling, we find a unique transcriptomic signature indicating mobilization of erythroid differentiation, a known response to cellular stress in this cell line (Figure 5C&D).

Treatment of control K562 with an ATR inhibitor induced changes in gene expression, although not on the scale observed with genetic perturbation of the TIP60 complex (either BRD8 or DMAP1). However, many of the genes found to be differentially expressed upon ATRi treatment are shared with the TIP60 perturbed cells and again indicate activation of a transcriptional network associated with differentiation. In the subset of genes whose expression changes exclusively upon ATRi we observed an upregulation of key cell cycle markers under direct influence of the S-phase checkpoint, such as CDC25A. Importantly, we find in the sgBRD8 exclusive genes a clear downregulation of apoptotic factors such as NFKBIA and numerous metabolic genes such as those involved in fatty acid oxidation and upregulation core histones.

Finally, we compared the effect of perturbing ATR in sgBRD8 cells relative to control sgBRD8 cells. We found no significant differential expression of any genes in this comparison (Figure 5E-H). These results suggest that loss of activity of the TIP60 complex may induce a cell state refractory to the effects of S-phase checkpoint inactivation. While surprising that a genetic perturbation could yield cells entirely unsensitized to drugging of an essential cell cycle regulator, this transcriptomic data agrees with the protective effect of sgBRD8 seen in our nominating and GI screen data. These findings support the notion that genetic rewiring (dGIs) can only be captured by comparative analysis of GI and eGI maps and would not be revealed through analysis of single gene perturbations, single drug perturbations or single drug/gene combinations using either single cell or population level readouts.

### Environmental GI maps in etoposide and glucose deprivation illustrate coordinating ontologies that drive rewiring

Our findings on the conditional relationships of the TIP60 complex in the context of ATR inhibition made us consider the extent to which the trend of a singular ontologically consistent set of genes driving large changes in functional relationships could be generalized to rewired interactions in response to other environmental conditions. To address this question, we performed three more GI mapping experiments in K562 cells using the same library of dual sgRNA targeting vectors. We created an additional reference (DMSO) GI map, an eGI map treated with the genotoxic chemotherapeutic, etoposide, and a third eGI map where the growth media used for the duration of the experiment was not supplemented with glucose. Etoposide is an effective toxin in replicating cells due to its inhibition and trapping of topoisomerase II on DNA, leading to replication fork blockage and double strand break formation (*43*). We reasoned that the cellular response to etoposide requires many of the same DNA damage and cell cycle regulators as for ATR inhibition, but the direct nature of the DNA damage induced may shed light on differences between these perturbations. We also chose to create a glucose deprivation eGI map as this perturbation represents an environmental nutrient perturbation that is quite distinct from the ATRi and etoposide perturbations. We reasoned this might be interesting because many genes related to metabolism and energy production are present in the library. As before, all three new maps were generated as two independent biological replicates (GI and eGI matrices for reference, etoposide treated, and glucose deprived experiments: Data S6, S7, and S8, respectively).

Replicate correlations at the sgRNA and gene level were similar to those observed in the first two GI experiments (Pearson R = 0.63, 0.64, and 0.55 for DMSO, etoposide, and glucose deprivation gene level, respectively. Figure S5A-C). After setting thresholds for significance at four times the standard deviation of the gene-ntc distribution, as done for previous GI and eGI maps, we find 191 positive and 548 negative interactions in the reference DMSO GI map, 774 positive and 698 negative interactions in the etoposide eGI map, and 187 positive and 434 negative interactions in the glucose deprivation eGI map (Figure S6A-C).

Gene and replicate averaged GI scores from the first reference map correlate highly with the same measurements paired from the second reference map (Pearson R =0.69, Figure 6A&B). In support of our methodology for measuring rewired genetic interactions, the dGI signal measured between paired reference maps is very low and weak, with only 22 total gene pairs exhibiting a false dGI, indicating that strong and/or ontologically consistent dGI measurements are indicative of real distinctions in functional relationships across conditions.

**Figure 6.**
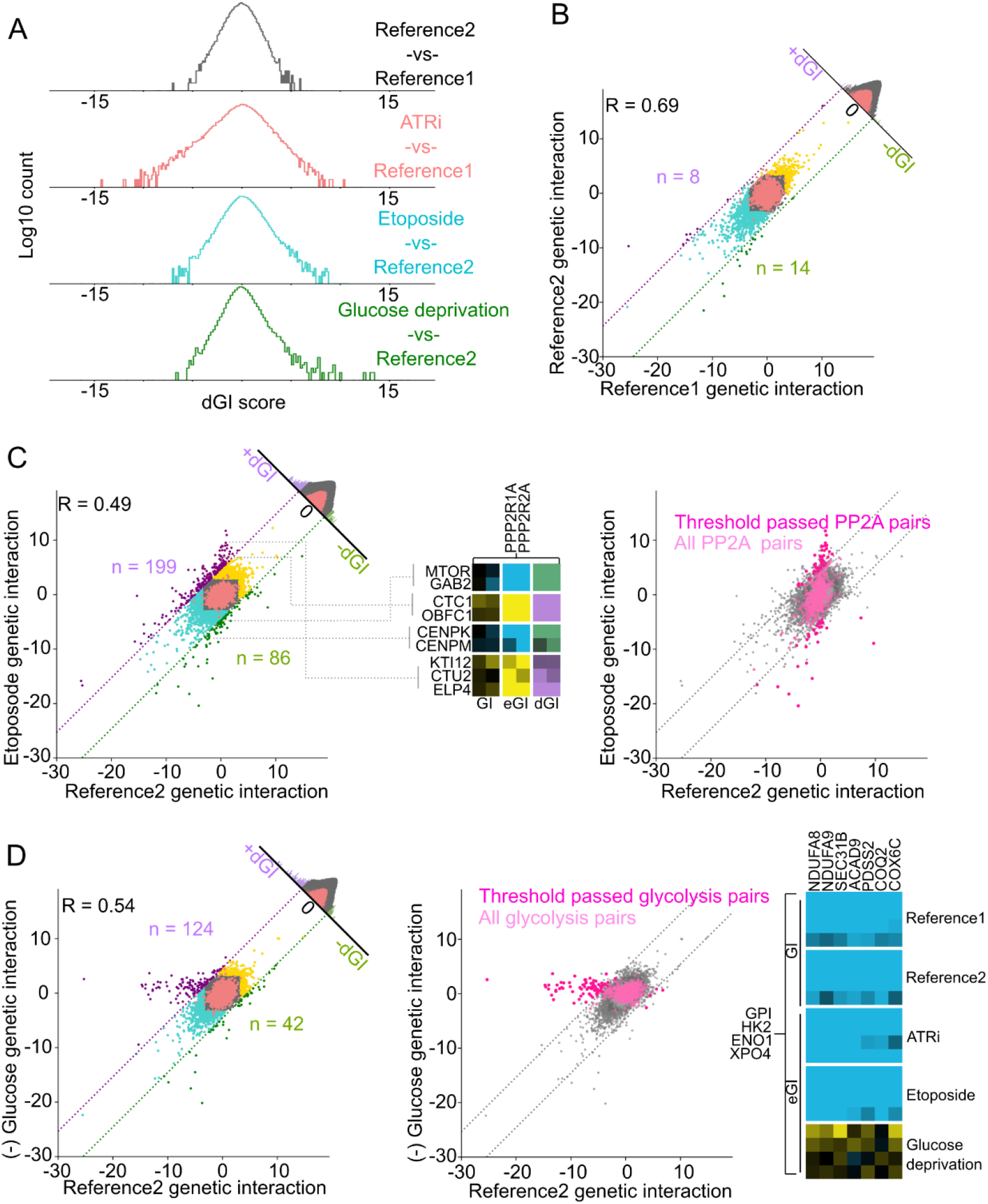
GI and eGI maps for reference, etoposide treatment, and glucose deprivation. **(A)** Distribution of dGI scores for reference, ATRi, etoposide treated, and glucose deprivation experiments (black, red, blue, and green, respectively). **(B)** Scatterplot of GI scores from both reference maps. Histogram on the diagonal displays dGI scores for this comparison, compare to Figure 4B. Regions of the plot shaded blue or yellow denote synthetic sick or buffering interactions, respectively. Gene-ntc pairs are highlighted in red. **(C)** Left) Same comparison as (B) but for etoposide treated eGIs versus paired reference GIs. Middle) Relationships between PP2A genes and example genes representing conditionally interacting ontologies. Right) All PP2A GI scores highlighted in pink with threshold passed dGIs in darker pink. **(D)** Left) Same comparison as (B) but for glucose deprived eGIs versus paired reference GIs, Middle) All glycolysis GI scores highlighted in pink with threshold passed dGIs in darker pink. Right) Relationships between glycolysis and oxidative phosphorylation gene sets for both reference and all three environmental GI maps.

Gene level and replicate averaged interactions measured in the etoposide and glucose deprivation eGI maps both correlate well with GI scores from the second reference map (Pearson R = 0.49 and 0.54, respectively, Figure 6C&D). However, as seen before, substantial numbers of interactions are specific to either the reference or environmental maps, suggesting rewiring of functional connections and consequently, differential genetic interactions. We created two new dGI matrices by taking the difference of eGI scores and matched reference GI scores for both the etoposide and glucose deprivation maps (Figure 6A and Data S9&S10). Differential genetic interactions measured from the etoposide map highlight gene sets with known rolls in replication fork fidelity, such as the CST complex, which protects telomeres and stalled replication forks from aberrant exonuclease activity (*44*). In the reference map, the CST complex is found to negatively interact with the BRCA1 complex as well as Replication Factor C (RFC) genes, which are critical for the loading of the Proliferating Cell Nuclear Antigen (PCNA). However, in the etoposide treated map, the relationships between these sets of genes are disrupted. Furthermore, the CST complex’s clustering is altered, as it exhibits an eGI profile that highly correlates with Non-Homologous End Joining (NHEJ) factors such as TP53BP1, SHLD1, and SHLD2.

Beyond the specific examples above, we also observed ontology level rewiring highlighted by the PP2A complex. In response to treatment with etoposide, the PP2A complex forms strong pervasive positive eGIs with a number of ontologies important for DNA replication, fork stabilization, spindle assembly, and translation regulation (Figure 6C). In every case, we find that the strongly deleterious effects resulting from loss of activity of these ontologies in the presence of etoposide are mitigated by the simultaneous perturbation of PP2A. This coordinating behavior of the PP2A complex, where environmental challenge broadly alters sets of functional connections for a single ontology, mirrors trends in our observations with TIP60 after ATR inhibition. While it is known that PP2A is required for mediating myriad distinct cellular processes, the patterns we observe suggest that environmental context significantly modifies its function. Specifically, our data suggests that PP2A is responsible for mediating the cytotoxicity of etoposide, perhaps indicating that PP2A is a key player in initiating p53 independent apoptosis when replicative damage exceeds the cell’s capacity for repair.

In contrast to the ATRi and etoposide maps, the resulting eGI scores from the glucose deprived cells correlated with those from the matched reference control to a high degree (Figure 6A&D). However, notable in this comparison is a set of interactions with very strong effect sizes in the reference GI map that are entirely missing from the glucose deprivation eGI map. Examining the identity of these highly conditional interactions reveal that nearly all of them include at least one gene necessary for glycolysis, and the strongest conditional interactions all derive from pairs between glycolytic genes and those important for oxidative phosphorylation or mitochondrial homeostasis. In both reference maps, the ATRi eGI map, and the etoposide eGI map, this set of interactions are consistently and uniformly among the strongest negative relationships measured in this study. However, this powerful and highly reproducible effect is entirely eliminated when cells are deprived of glucose in their growth media (Figure 6D). We believe this conditionality derives from environmental epistasis to a genetic perturbation - the removal of glucose induces a cell state where glycolytic genes are functionally inactive, severing their relationships with other metabolic processes. The glucose deprivation eGI map thus yields the most specific and targeted set of dGIs we’ve measured: other than the rewiring between metabolic factors, nearly all interactions are consistent between the environmental and reference maps. In summary, comparison of the three dGI maps demonstrates that in each environmental context, profiles of dGIs are highly context specific and are frequently found among genes that have an annotated function that relates to the perturbation in question.

### Integrated analysis of GI and eGI matrix clusters enables higher level analysis of ontology level rewiring

Initial inspection of our five maps showed that hierarchical clustering patterns within each reference and environmental GI map revealed that ontologically consistent clusters rarely exhibited changes in gene lists within the cluster. To more rigorously evaluate intra- and inter- complex correlations (and thus clustering) across maps, we were interested if the breadth of GI and eGI profiles collected across the five maps of this study could be integrated in order to form a consensus clustering that would serve to enable more robust comparisons. To achieve this, all five maps were independently normalized by their standard deviation, in order to mitigate jackpotting by the particularly strong signal measured in the ATRi eGI map. Next, all maps were concatenated on one axis to create a “n” by “5n” matrix of all interactions measured. Finally, the short end of the consensus matrix was hierarchically clustered using the average pairwise linkage of the Pearson correlation. The distance matrix of this clustering could then be applied to each individual map in turn in order to form a standardized set of ontologies for the entire study (Figure 7A).

**Figure 7.**
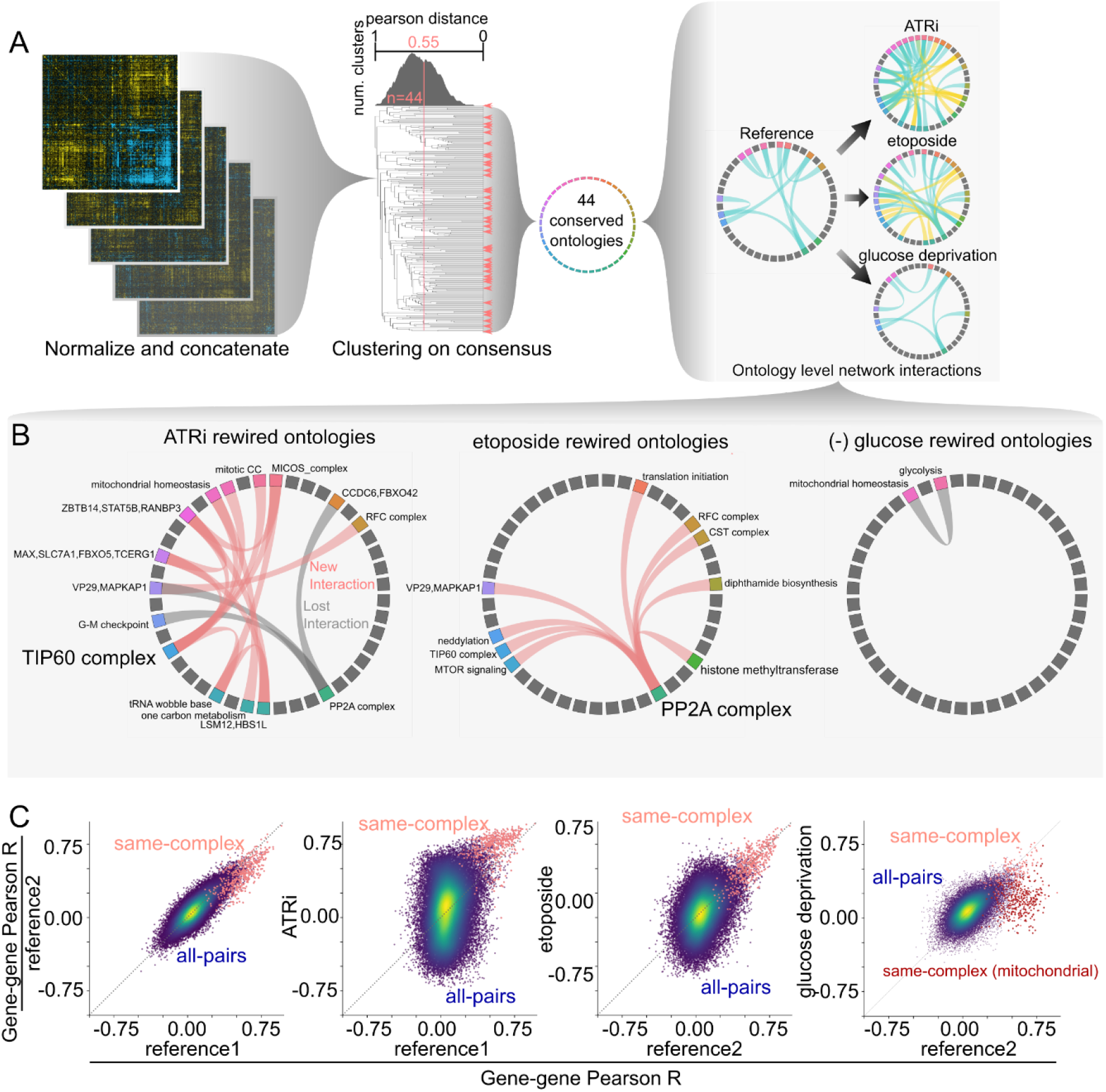
Ontology level measurements of environmentally induced functional rewiring. **(A)** Schematic of discovering conserved ontologies by consensus clustering of two reference GI and three eGI maps. Left) Each GI map was normalized to its respective standard deviation, then concatenated to form a consensus matrix. Middle) This matrix is clustered using the average pairwise linkage of the Pearson correlation, as before. A Pearson distance threshold is applied that maximizes cluster formation without collapsing discrete ontologies, yielding 44 conserved gene lists (see methods). This analysis can be used to generate ontology level genetic interaction maps as a method of visualizing systems level information. **(B)** Systems level rewiring of complexes and ontologies. Inter-ontology relationships that are specific to the given environment are linked with pink edges (new interactions). Relationships that are found in the reference condition but not the environment are shown in gray (lost interaction). **(C)** Scatterplot of all gene-gene GI profile correlations between two reference GI maps or a reference and environmental GI map. Gene pairs where both belong to the same conserved cluster as defined in Figure 7A are highlighted in red. Dark red points denoting gene pairs belonging to the mitochondrial homeostasis cluster are highlighted in comparison for the glucose deprivation map versus the second reference map.

In order to validate that this consensus method did not corrupt clustering information, we applied the framework above to the sgRNA level GIs from each map. Most genes in this experiment are represented by two independent sgRNAs so the correlation between same-gene sgRNA profiles can be used as a quantitative metric of the clustering fidelity of a dataset. We find that the median Pearson correlation for same-gene sgRNA profiles in the consensus sgRNA matrix is 0.37. This score is consistent with or better than the same metric in all independent sgRNA matrices with the exception of the ATRi eGI sgRNA matrix, which is modestly higher (Figure S6D-I).

After applying our consensus clustering framework to the five gene level maps, we generated a list of ontologies in our data and the consensus set of genes that belong to them. We established a threshold in the distance matrix at a Pearson correlation of 0.55, as this maximizes the number of clusters without grouping together sub clusters, which could result in information loss (Figure S6J). Genesets from each of these 44 clusters were manually tested against the Molecular Signatures database (MSigDB) in order to determine if there exists a known ontology associated with them. MSigDB finds 29 out of 44 consensus clusters have at least one strongly (p-value 1.0e-5) associated GO term in its “Biological Process” or “Molecular Function” datasets (Data S11).

Consensus clusters with no predicted gene ontology association are interesting as they implicate a set of genes with no known relationship to have a conserved function. PRPF39 and TRNAU1AP are two such genes that have predicted RNA binding and processing domains and conserved clustering but no established function. Not only do these genes associate with one another, they also form a close and conserved relationship with CBLL1, ZC3H13, and KIAA1429, three genes known to belong to the WMM complex, which is crucial for adenosine methylation in mRNA maturation. This association suggests that PRPF39 and TRNAU1AP might be unidentified members of the WMM complex or otherwise coordinate their predicted RNA processing activities to the goal of mRNA maturation. Such examples illustrate the power of integrated GI experiments to group known and unknown genes together and generate hypotheses.

For the 44 sets of genes with conserved relationships we generated a compressed matrix for each of the five GI experiments by averaging the GI or eGI score of all possible gene pairs within those clusters. As with gene level scores, we then took the difference between each environmental ontology matrix and its matched reference matrix to form a differential ontology matrix (Figure 7B). This data illustrates the degree of rewiring in each environment at a higher functional level of complexity relative to gene level analysis. Circos plots of these data provide an immediately interpretable summation of differential genetic interactions and a strong visualization of the coordinating ontology behavior exhibited by the TIP60 complex, PP2A complex, and glycolysis in the ATRi, etoposide, and glucose deprivation dGI maps, respectively.

Finally, we compared profiles of gene level Pearson correlations between experiments as a measure of the conservation of clustering information across conditions (Figure 7C). The two reference experiments are highly correlated while all environment-reference comparisons display varying degrees of correlation. In particular the ATRi and etoposide maps exhibit an increasing cooperation between genes involved in disparate bioprocesses, as indicated by exaggerated positive correlation distributions relative to other maps. Conversely, the ATRi and etoposide maps also exhibit negative correlation distributions relative to other maps suggesting dissociation of genetic relationship. Perhaps the genome destabilizing effects of these drugs require closer associations and also different associations between various functional modules to allay cell death, which would be reflected in broad correlation distributions. For the most part, gene pairs belonging to the same protein complex are found to correlate highly with one another in each comparison. However, this trend is not conserved for the glucose deprivation map, where disruption of interactions within the mitochondrial supercluster drives a relatively low level of clustering conservation. In general, our data illustrates that relationships between same-complex genes are broadly conserved regardless of environmental challenge, while inter-ontology associations prove to be significantly more plastic. Together these data and analytic tools provide a compelling case for extending the intellectual framework of genetic interaction mapping to measure the plasticity of genetic relationships in respect to environmental challenges.

## Discussion

Here, we show that genetic interaction network connectivity is functionally rewired in human cells exposed to different environmental conditions using cell cycle interruption, genotoxic perturbation, and nutrient deprivation as archetypes. To our knowledge, this study constitutes the first systematic measurement of context-specific genetic interactions under different environmental states in human cells (*5*, *10*, *22*, *23*, *26*, *27*). Our study was enabled by our development of a scalable and quantitative GxGxE GI mapping platform along with new analytic strategies, both of which are required for measuring and interpreting rewired GIs. We find that while intra-complex genetic interactions tend to remain consistent across environmental exposures, each environmental condition exhibits specific changes to inter-complex/inter-pathway connections that reveal unexpected conditional genetic dependences and suggest new mechanistic hypotheses for future study.

Our observation that each set of rewired genetic interactions are environmentally context-dependent and primarily driven by small numbers of unexpected key genes, raises an intriguing concept that there could be a very large number of specific ways in which gene networks have evolved to respond to environmental perturbations (*45*). For example, in the context of S-phase checkpoint biology, we identify the TIP60 histone acetyltransferase complex as a driving force for dGIs in the ATR inhibited genetic landscape. TIP60 displays rewired GIs with many ontologies in the map, particularly those involved in cell cycle modulation, gene expression, and metabolism. Knockdown of TIP60 leads to large changes in gene expression and, critically, results in a transcriptional cell state that is seemingly epistatic to ATR inhibition. Similarly, upon treatment with etoposide which activates DNA damage, we observed ontology level rewiring between the PP2A complex and a number of ontologies important for DNA replication, fork stabilization, spindle assembly, and translation regulation which suggests that environmental context significantly modifies PP2A function in cellular homeostasis. Lastly, in the context of glucose withdrawal, we observe that deprivation of glycolytic enzyme substrates results in a severing of the functional connections between glycolytic genes and other metabolic genes suggesting the cellular environment can rapidly modulate functional connections governing metabolic flux.

In summary, our findings are among the first systematic pieces of evidence to demonstrate models that imply a one gene, one function relationship is likely not sufficient to fully explain complex features of human cell biology and reveal the surprising degree to which gene relationships are plastic.

## Acknowledgements

Our thanks to April Pawluk, Jason Swinderman, Chris Hsiung, Caroline Wilson, and Arc SciPubs for reviewing the manuscript, and consulting on data presentation, analytic strategy, and clarity. Sequencing was performed at the UCSF CAT, supported by UCSF PBBR, RRP IMIA, and NIH 1S10OD028511-01 grants.

## Author Contributions

B.W.H. and L.A.G. were responsible for planning experimental design. B.W.H. performed all experiments described and prepared and cloned all new reagents required. B.W.H., G.T.W., and T.M.N. were responsible for data analysis, statistics, and computational approaches. B.W.H. wrote and assembled the manuscript. L.A.G., T.M.N., and G.T.W. critically edited the manuscript.

---

**Figure S1.**
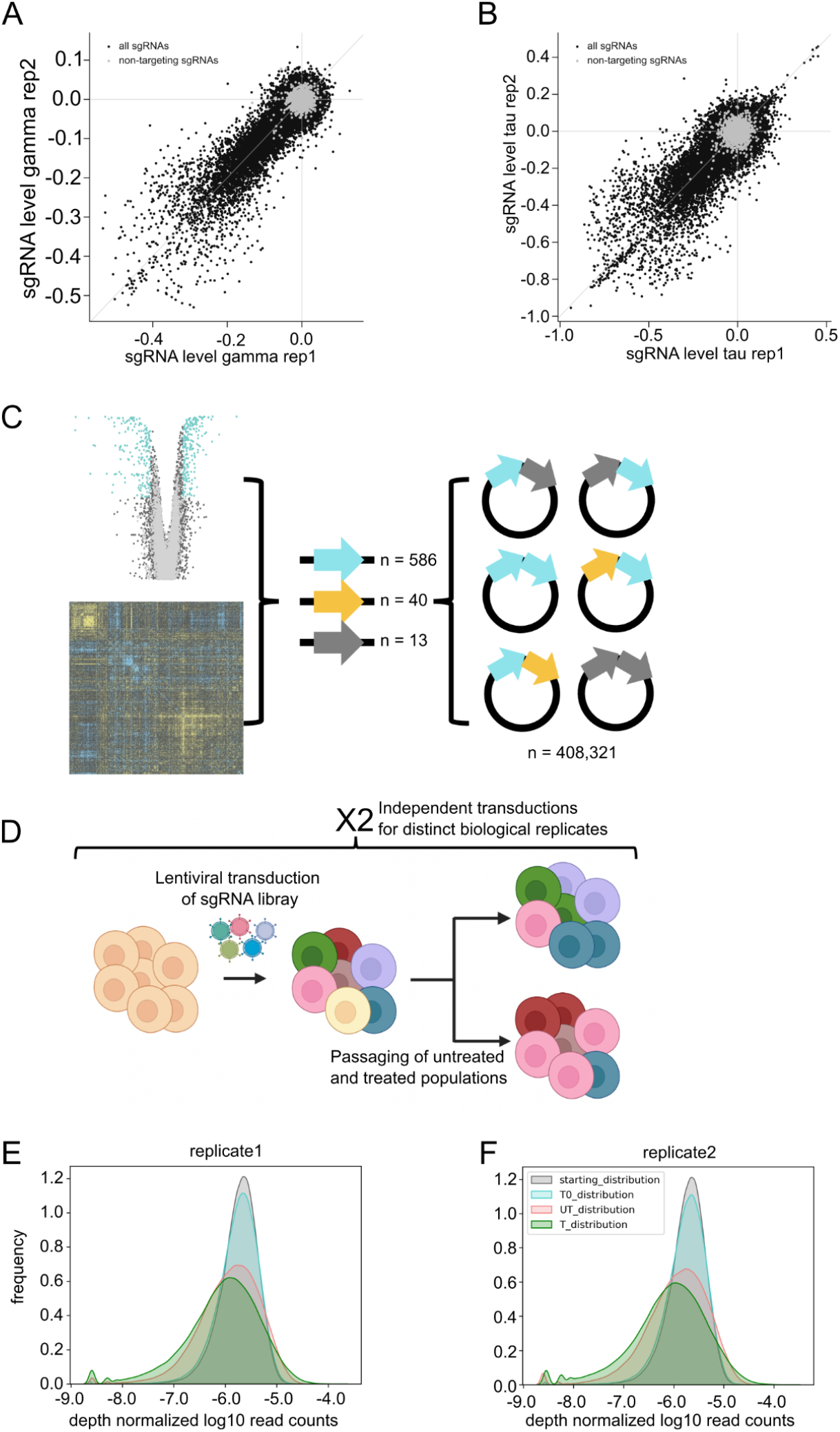
Nominating screen and paired library cloning quality control. **(A&B)** Scatterplots of sgRNA level phenotypes from the control (A) and ATRi treated (B) arms of the genome-scale CRISPRi nominating screen (gamma and tau scores, respectively, see methods). Each individual gene targeting sgRNA shown in black. Non-targeting control sgRNAs are shown in light gray. **(C)** A schematic of our GI mapping dual-sgRNA library design and sgRNA inclusion criteria. Cyan arrows denote sgRNAs that induce an ATRi specific growth response validated from our ATRi nominating screen. Yellow arrows denote sgRNAs validated from previous studies (*20*) that are relevant to DNA repair and cell cycle biology but not included from the nominating screen. Gray arrows denote non-targeting control sgRNAs picked from the nominating screen based on their lack of phenotype. **(D)** Schematic for screening in both the nominating CRISPRi and GI screens in K562 cells (see methods for details) **(E&F)** Kernel density plots of read counts mapped from dual-sgRNA GI library for both independent replicates normalized by the sum of reads in each condition shown: pre-transduction, timepoint zero, final timepoint control, final timepoint ATRi treated (gray, blue, red, and green, respectively).

**Figure S2.**
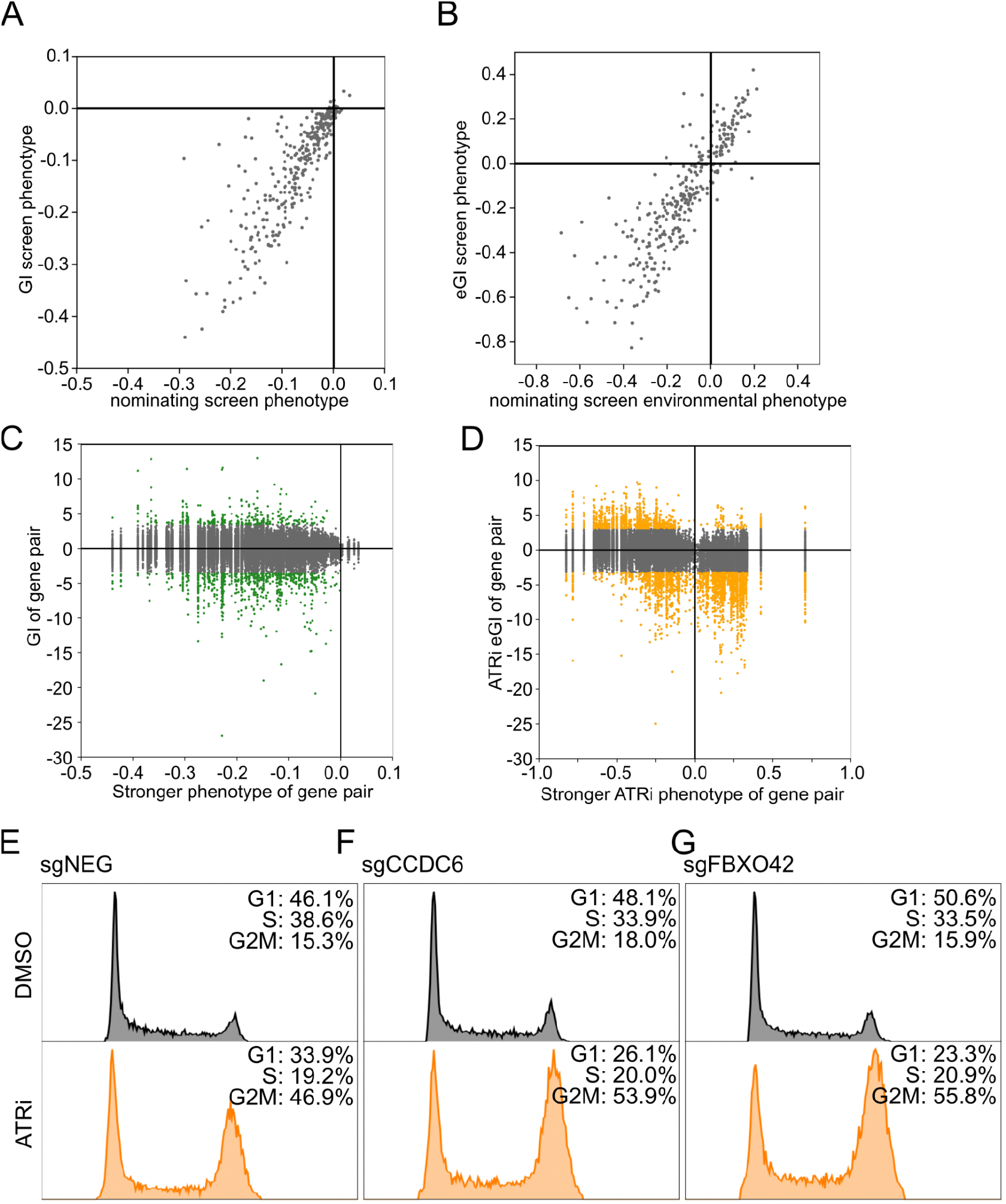
GI map quality control and phenotype validation. **(A&B)** Scatterplots of the gene level control or ATRi treated phenotypes (gamma and tau, respectively) calculated in our nominating genome-scale CRISPRi screen versus the same phenotypes calculated in our GI and eGI screens. **(C&D)** Scatterplots of the stronger (farthest from zero) sgRNA control (C) or ATRi treated (D) phenotype calculated for each pair of sgRNAs versus the GI or eGI calculated between that pair. Gene pairs with a significant GI or eGI score are highlighted in green and orange, respectively. **(E-G)** Propidium iodide staining for DNA context of dividing cells expressing a non-targeting sgRNA (E), sgCCDC6 (F) or sgFBXO42 (G) under basal (DMSO) or ATR inhibited conditions.

**Figure S3.**
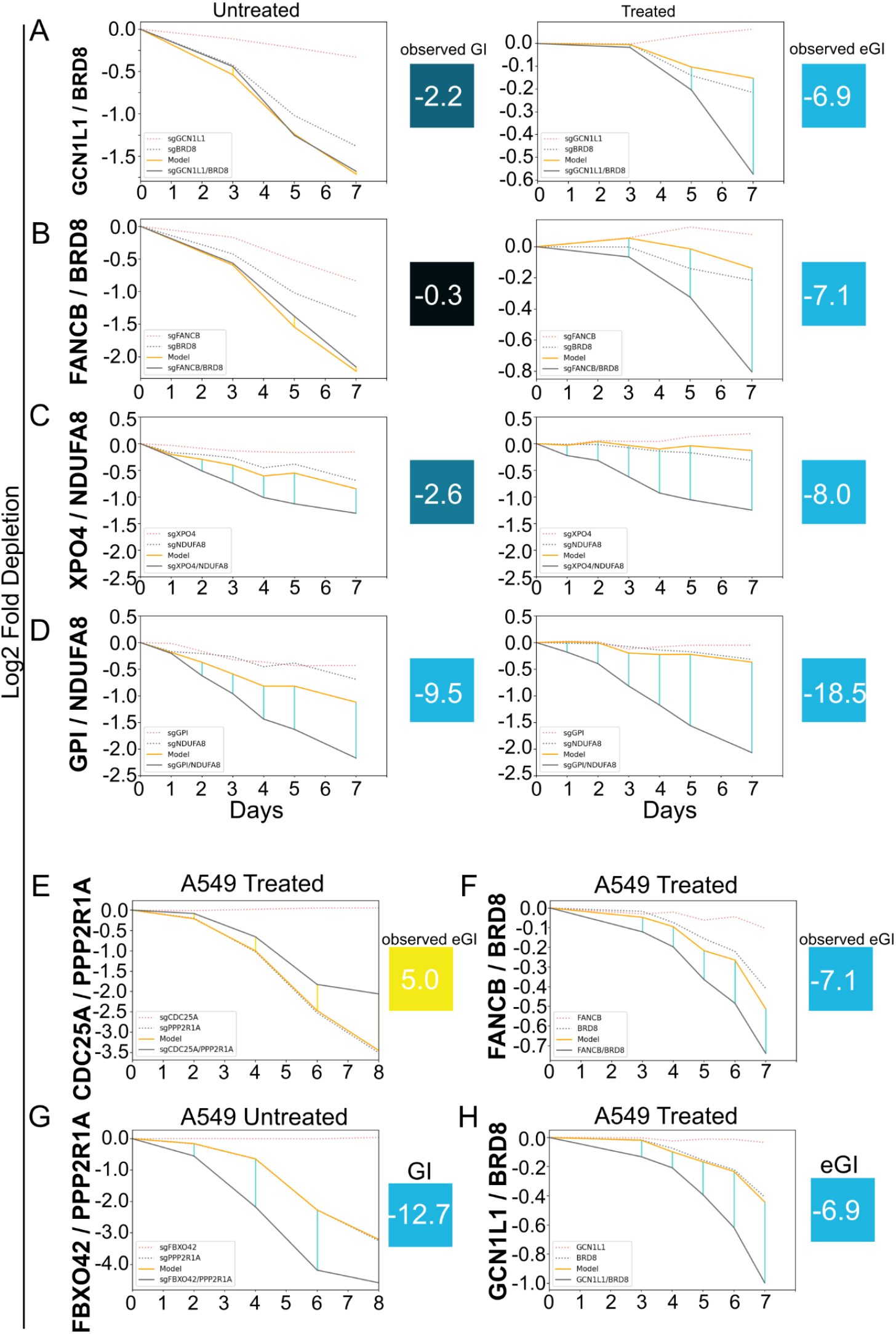
Fluorescence competition assays. **(A-D)** Assays to validate environmental conditionality of interactions. Depletion phenotypes of single gene perturbations over time are shown as dotted lines. Modeled paired phenotypes are shown as orange solid lines. Observed paired phenotypes are shown as gray solid lines. Interaction scores are shown as vertical lines between modeled and observed paired phenotypes at all timepoints measured, either blue or yellow to denote a negative or positive interaction, respectively (see methods). The GI and eGI scores shown to the right of the plots are the observed interactions from our maps. **(E-H)** Assays to validate interactions in A549 cells. Plot structure is the same as in A-D.

**Figure S4.**
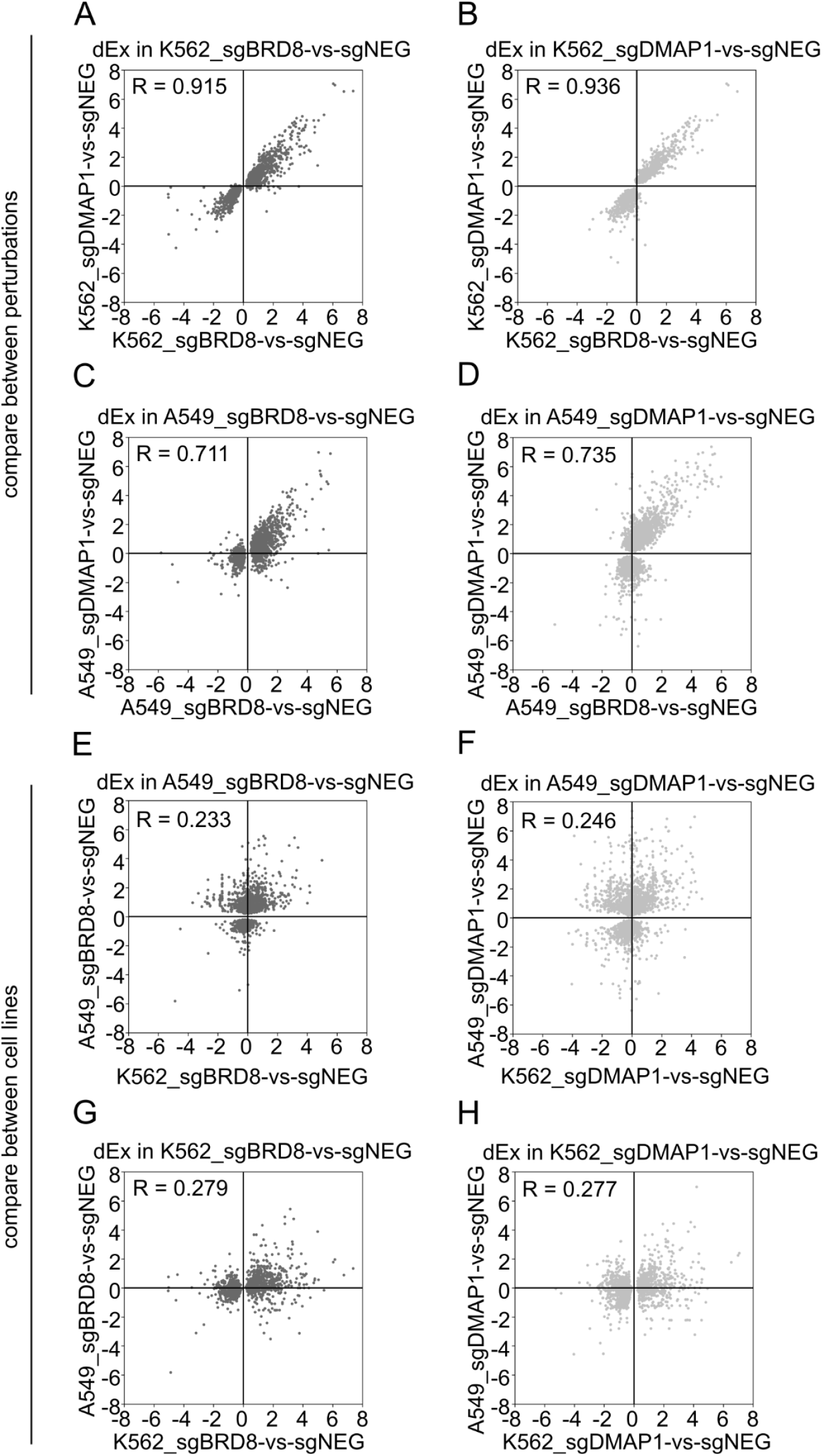
Comparisons of differential expression values across perturbations and cell lines. (A-H) Scatterplots of differential expression (dEx) values calculated using DEseq2. Each point corresponds to a gene in a set of genes that is found to be significantly differentially expressed in the comparison in the header of each plot. A Pearson R value for the comparison of each differential expression comparison is listed in the top left of each plot.

**Figure S5.**
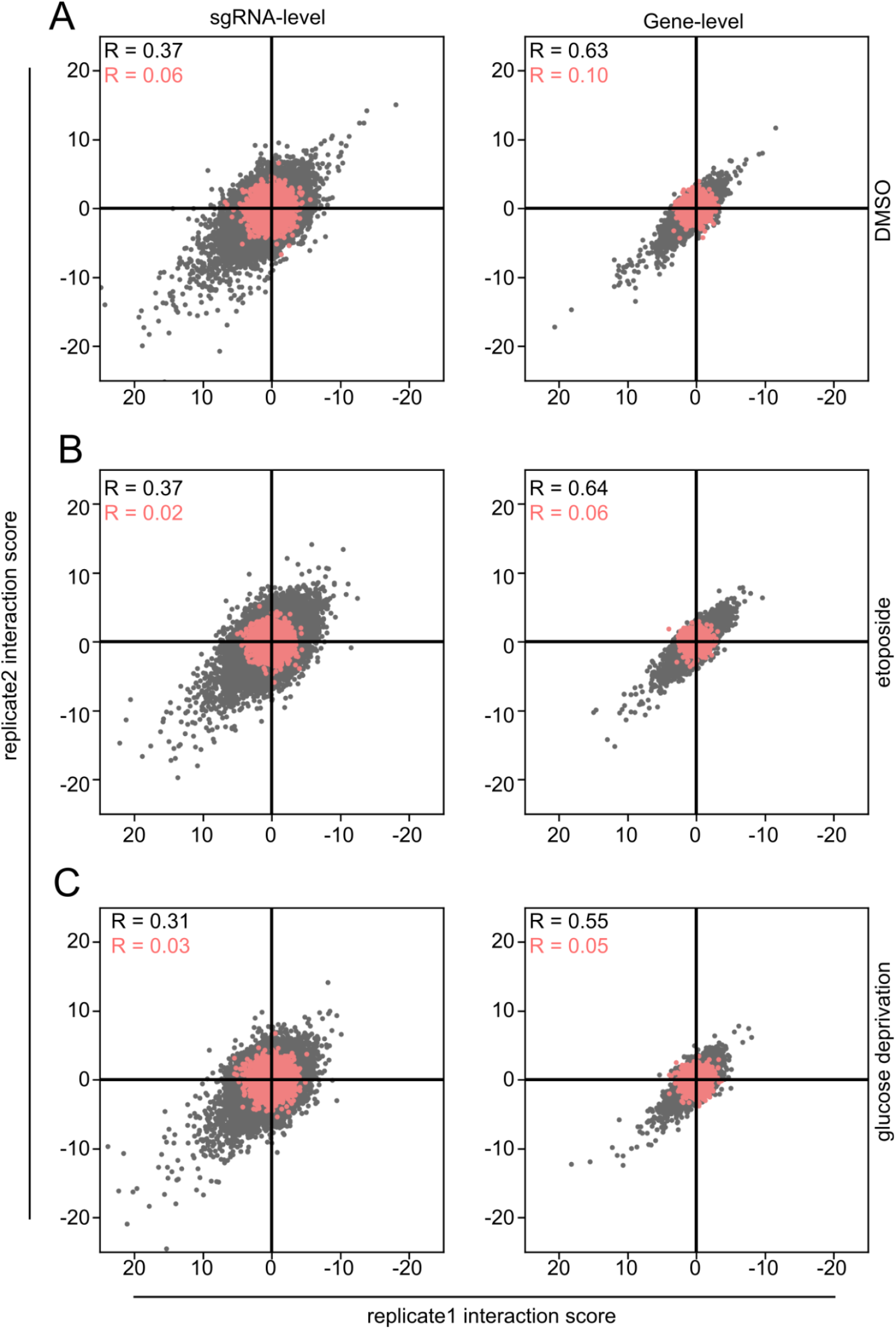
Replicate quality control of GI and eGI experiments. **(A-C)** Scatterplot of sgRNA (left column) and gene (right column) level interaction scores for DMSO (A), etoposide treated (B), or glucose deprived (C) GI and eGI maps. Genes or sgRNAs paired with negative control non-targeting sgRNAs shown in red. Pearson R values shown in top left of each plot.

**Figure S6.**
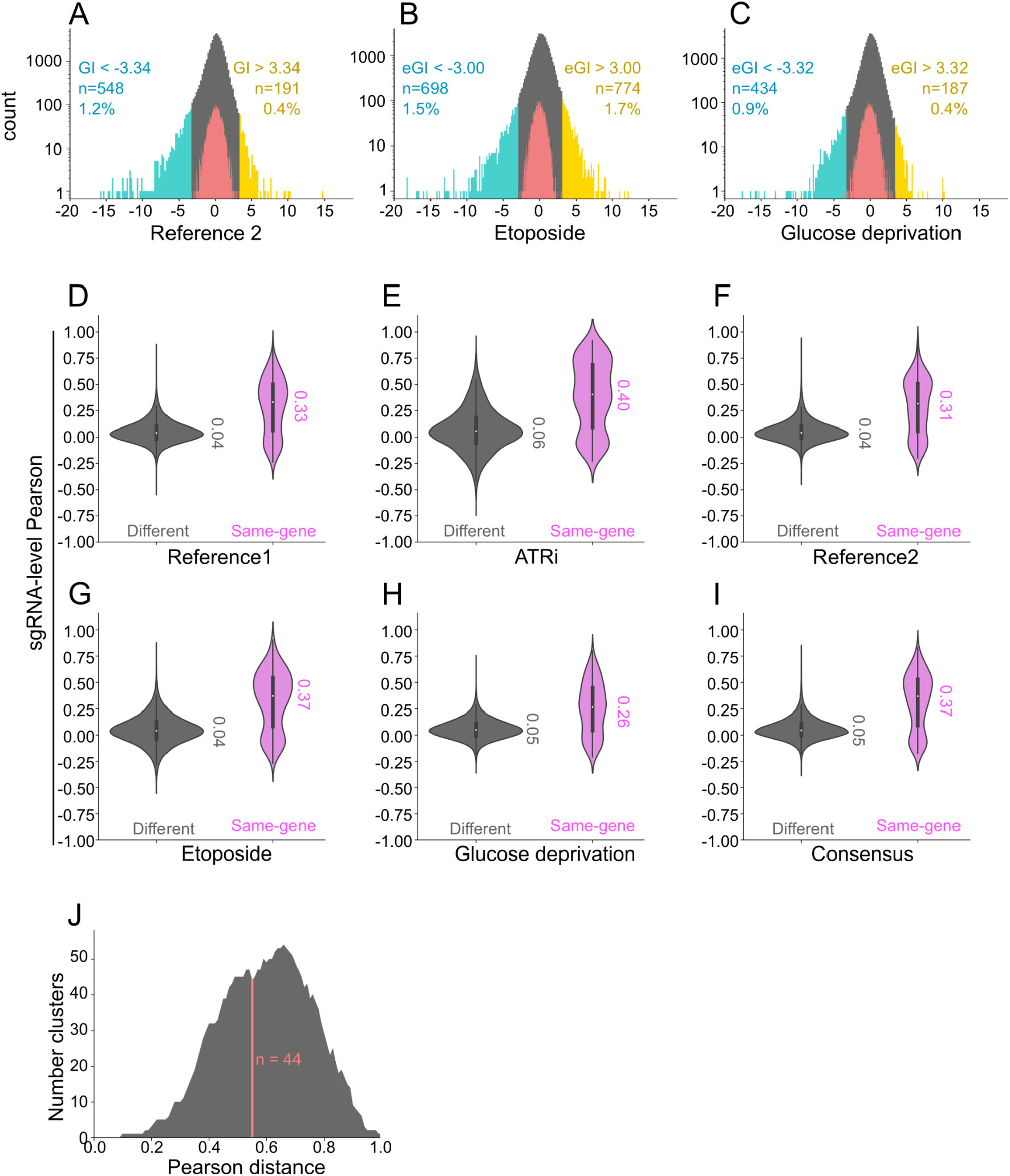
Interaction distributions and clustering quality of maps. **(A-C)** Interaction distributions for reference (DMSO), etoposide treated, and glucose deprived GI/eGI maps. Interaction score thresholds, number of threshold passed gene pairs, and percentage of total gene pairs measured displayed on both wings of all histograms. Significance thresholds are represented by histogram shading. Negative and positive interactions are highlighted in blue and yellow, respectively. Gene-ntc pairs are highlighted in red. **(D-H)** Violin plots of Pearson R distributions of sgRNA-level interaction profiles for all five GI/eGI maps. Gray violins denote sgRNAs that target different genes, purple violins denote sgRNAs that target the same gene. Median values for all distributions displayed to the right of each violin. **(I)** Same comparison as in D-H but for the consensus matrix described in Figure 7A. **(J)** Number of clusters formed from the consensus matrix data as a function of Pearson distance. Line in red shows cluster formation at a Pearson distance threshold of 0.55.

**Data S1. Nominating genome scale ATR inhibitor CRIPRi screen results.** Gene level growth scores and associated p-values for untreated perturbation phenotypes (gamma), ATRi treated perturbation phenotypes (tau), and treated normalized by untreated perturbation phenotypes (rho) (see methods). Genes with a |rho| score >= 0.2 and rho p-value <0.02 were candidates for inclusion in our GI library.

**Data S2. Validated single sgRNA list used in construction of paired sgRNA GI library.** Includes gene targets and protospacer sequences. All sgRNAs have been validated as active either in our nominating screen (Figure 1, Data S1) or in a previously published study (*20*). Negative control non-targeting sgRNAs (ntc) were selected from the pool of pseudogenes in our nominating screen with phenotypes closest to zero.

**Data S3. Matrix of reference experiment 1 GI scores.** All pairwise genetic interactions between genes in our GI library for the first set of reference (DMSO) experiments

**Data S4. Matrix of ATRi environmental GI scores.** All pairwise ATRi environmental genetic interactions between genes in our GI library.

**Data S5. Matrix of differential ATRi GI scores.** All pairwise differential ATRi genetic interactions between genes in our GI library. Calculated by taking the difference of each gene pair’s ATRi eGI and reference map 1 GI scores.

**Data S6. Matrix of reference experiment 2 GI scores.** All pairwise genetic interactions between genes in our GI library for the second set of reference (DMSO) experiments

**Data S7. Matrix of etoposide environmental GI scores.** All pairwise etoposide genetic interactions between genes in our GI library.

**Data S8. Matrix of etoposide environmental GI scores.** All pairwise glucose deprivation genetic interactions between genes in our GI library.

**Data S9. Matrix of differential etoposide GI scores.** All pairwise differential etoposide genetic interactions between genes in our GI library. Calculated by taking the difference of each gene pair’s etoposide eGI and reference map 2 GI scores.

**Data S10. Matrix of differential glucose GI scores.** All pairwise differential glucose genetic interactions between genes in our GI library. Calculated by taking the difference of each gene pair’s glucose eGI and reference map 2 GI scores.

**Data S11. List of conserved ontologies from consensus clustering.** A list of all sets of conserved ontologies derived from our consensus matrix under a Pearson distance threshold of 0.55. Clusters are annotated with MSigDB if at least one term in the “Biological Process” or “Molecular Function” datasets is enriched with a p-value less than 1.0e-5. If no GO term is found sufficiently enriched, the cluster is labeled “no_GO”. Gene sets for each cluster are listed.

## Experimental Materials & Methods

### Mammalian cell line culture and lentivirus preparation

All mammalian cell lines were cultured at 37°C with 5% CO2. K562 cells stably expressing dCas9-KRAB (*17*) were grown in RPMI-1640 media with 25mM HEPES (GIBCO), supplemented with 10% FBS, 100units/mL Penicillin, 100µg/mL Streptomycin, and 292µg/mL Glutamine (GIBCO). During the nominating and GI screens, K562 were grown in shaker flasks in a dedicated cell culture shaker/incubator (INFORS) at 120rpm and the media these cells were grown in was further supplemented with 0.1% Pluronic surfactant (GIBCO). A549 cells stably expressing dCas9-KRAB were grown in F12K media supplemented with 10% FBS, 100units/mL Penicillin, 100µg/mL Streptomycin, 292µg/mL Glutamine (GIBCO), 1x MEM Non-Essential amino acids (GIBCO), and 1mM Sodium Pyruvate (GIBCO). HEK293T cells were grown in DMEM media supplemented with 4.5g/L D-Glucose, 110mg/L Sodium Pyruvate (GIBCO), 10% FBS, 100units/mL Penicillin, 100µg/mL Streptomycin, 292µg/mL Glutamine (GIBCO). Lentivirus was prepared from HEK293T cells using TransIT-LT1 transfection reagent (MIRUS) and standard three-vector packaging plasmids. 72hr post-transfection, viral supernatant is filtered with 0.45µm filters and frozen at -80°C.

### sgRNA expression vector and library design

All sgRNAs used in all experiments were derived from a previously described list of the top5 algorithmically determined CRISPRi guides for each protein-coding gene in the genome (*29*). This genome-scale CRISPRi library was used for the nominating CRISPRi ATRi screen.

To clone the library used for GI mapping: the two sgRNAs with the strongest phenotype from each of the 293 genes in the nominating screen with a |**ρ**| score (see below) >0.2 and -log_10_ Mann-Whitney p-value >2.0 were used as the basis for our focused GI library. To this list of 586 sgRNAs, 40 more were included targeting 20 genes with a known connection to a range of gene ontologies pertaining to the DNA damage response that had been validated in a previous study (*20*). Finally, 13 negative control sgRNAs were included bringing the total size of the library to 639 unique sgRNAs (Figure 2B, Data S2). Protospacers for this library were obtained in a pooled format commercially, subsequently cloned into two independent intermediate vectors, then cloned into the final dual sgRNA expression vector in which every possible pairwise permutation of sgRNAs in the singles library is represented, as described previously (*20*). Through attrition inherent in cloning large sgRNA libraries, a small percentage (<1%) of sgRNA pairs were lost, resulting in a realized library size of 405,667 with 90% of all elements expressed at <16-fold variation (Figure S1E&F).

Dual sgRNA vectors used in fluorescence competition validation experiments were cloned in the same expression vector as used in the GI library, in an arrayed format.

Single sgRNA vectors used in RNAseq experiments were cloned into the same vector used in the nominating CRISPRi screen, in an arrayed format.

### Transduction and screening with CRISPRi libraries

Transduction and passaging of cells was performed in duplicate as independent biological replicates for all experiments. For nominating and GI screens, K562 cells stably expressing dCas9-KRAB were transduced with a lentiviral preparation of an sgRNA library using 8µg/mL polybrene with the goal of obtaining a transduced population representing at least 250-fold coverage of the library while maintaining an MOI below 0.3. This population of cells is grown and expanded continuously and dosed with puromycin at a concentration of 1µg/mL 48 and 72 hours post transduction to select for infected cells. The infected population was expanded until at least 4000-fold coverage of infected cells were available to begin the screen. At the initial timepoint, cell pellets with 1000-fold coverage of the library were frozen in 10% DMSO freezing media. This initial timepoint was taken at day six and day seven post transduction for the nominating screen and all GI screens respectively. Final timepoint samples for all screens were collected as 1000-fold coverage of the library.

For the nominating screen, samples were split to a concentration of 2.5e5 cells/mL every other day in fresh media. For the ATRi treated arm of the screen, on days zero and seven the treated samples were dosed with 750nM AZD6738, and the untreated samples were dosed with an equivalent volume of DMSO. Fifteen days after the start of the experiment, once the cumulative difference in population doublings between the treated and untreated samples had approximately reached five, the final timepoint was taken.

For the ATRi GI screen, samples were split every day to a concentration of 5e5 cell/mL in fresh media. On days zero and seven of the screen, the treated sample was dosed with 1.25µM AZD6738, and the untreated samples were dosed with an equivalent volume of DMSO. Final timepoint samples were collected on day nine after the cumulative difference in population doublings between the treated and untreated populations exceeded 4.5 in at least one replicate.

For the etoposide GI screen, on day zero, the treated samples were dosed with 500nM etoposide and untreated samples with an equivalent volume of DMSO. Media was washed out and replaced with fresh untreated media 24 hours later. Three days after the start of the experiment, cells were again dosed with 500nM etoposide. On subsequent days cell populations were daily split back to a concentration of 5e5 cells/mL of media but without media washout, allowing a longer exposure time for the etoposide sample. Final timepoint samples were taken at day nine once differences in cumulative population doublings between the etoposide and control populations had reached 4.11 in at least one replicate.

For the glucose deprivation GI screen, on day zero, the treated samples were resuspended in RPMI media without glucose. Treated cells were grown in this type of media for the length of the experiment. Every day all samples were analyzed and split to a concentration of 5e5 cells/mL in fresh media. Final timepoint samples were collected on day nine once differences in cumulative population doublings between the glucose deprived and control populations had reached 2.33 in at least one replicate.

For both the nominating screen and all GI screens, genomic DNA was extracted from T0 and final timepoint cell pellets using a Macherey-Nagel Blood XL cleanup kit. Integrated sgRNA loci were amplified from gDNA using PCR to append adapters for NGS, then submitted for sequencing on an Illumina Hiseq4000 or NovaSeq platform, for the nominating and GI screens respectively. For GI libraries, a hamming distance of one is used when mapping reads to our reference library, to allow for minor errors in sequencing to not be excluded from analysis. Custom sequencing primers were used for both nominating screen and GI maps as described previously (*20, 29*).

### Fluorescence competition assays

K562 or A549 cells are transduced with a lentiviral preparation of a single or dual sgRNA expression vector and expanded continuously for several days. At day five post transduction, cells are seeded at a concentration of 5.0e5 cells/mL and dosed with 1.25µM AZD6738 or an equivalent volume of DMSO, constituting the d0 sample. Each condition is analyzed by flow cytometry everyday or every other day to determine the ratio of uninfected control cells to BFP+ transduced experimental cells before being reseeded at 5.0e5 cells/mL in fresh medium. This process is repeated until seven or eight days after the first dose or the transduced population falls below 1% of the total population.

Data is processed by normalizing all samples by their day 0 %BFP+ to calculate the deviation from the starting value. To account for random variation in data collection, each timepoint for each sample is then further normalized by the deviation observed in the ntc/ntc dual sgRNA sample at that same timepoint. The log2 of these values is taken to derive the enrichment/depletion phenotype. Next, a matrix of modeled dual sgRNA phenotypes is calculated by summing the phenotypes from samples with only a single targeting sgRNA at each timepoint. The difference between phenotypes from the observed dual sgRNA vectors and the modeled phenotype derived from the single sgRNA vectors is recorded as the validation genetic interaction.

### Propidium Iodide (PI) Staining of DNA content

Pure populations of cells transduced with an sgRNA targeting CCDC6, FBXO42, or ntc were grown in media treated with 1.2µM AZD6737 or an equivalent volume of DMSO for 24hr. Cells were washed and grown in fresh untreated media for another 24hr before harvesting for PI staining (total time post treatment = 48hr). To prepare for staining and flow cytometry, cells were washed in PBS (GIBCO) and cell pellets resuspended in ice-cold 70% EtOH while being vortexed. Fixation continued at -20°C for 1hr, then cells were washed in PBS and analyzed by flow cytometry to determine concentration. 5E5 fixed cells were spun down and resuspended in 500µL of PI/RNase staining solution (Invitrogen). Cells were stained for 30 minutes then analyzed by flow cytometry to determine DNA content distributions of each population.

### RNAseq

K562 cells are transduced with a lentiviral preparation of a single sgRNA expression vector targeting either BRD8, DMAP1, or ntc in replicate. 48hr post-transduction cell populations are sorted by %BFP+ to obtain a pure population of transduced cells and expanded continuously for several days. On day five post-transduction, cell populations are dosed with 1.5µM AZD6738 or an equivalent volume of DMSO. 48hr hours after dosing 1.0e6 cells are harvested from each sample. RNA is then extracted from cell pellets using a Zymo Direct-Zol kit. 350ng of RNA from each sample is used as the input for the Lexogen Quantseq FWD kit with UMI add-on. All steps of the kit are performed according to the manufacturer’s instructions.

### Analytic Methods

#### Calculating sgRNA level growth phenotypes from CRISPRi screens

Calculating primary sgRNA level growth phenotypes for both the nominating CRISPRi and GI screens were done in the same way. First, each sgRNA construct within a sample is normalized by the total read counts for all sgRNAs in that sample, to adjust for differences in sequencing depth. Phenotypes in an untreated condition (**γ**), or treated condition (**τ**) are calculated as:

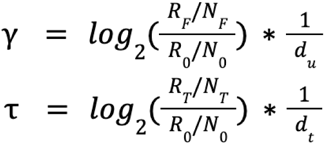

where **R_F_**, **R_T_**, and **R_0_**are the number of reads for the query sgRNA construct in the final untreated, final treated, and initial timepoints respectively. **N_F_**, **N_T_**, and **N_0_** are the median number of reads among negative controls (ntc-ntc pairs in GI screens) in the final untreated, final treated, and initial timepoints, respectively. **d_u_** and **d_t_**are the cumulative number of cell population doublings in the experiment in the untreated and treated conditions, respectively.

Additionally, to calculate the guide normalized effect on cell viability in the treated condition, that is used to threshold hits in the nominating screen (**ρ**), we use:

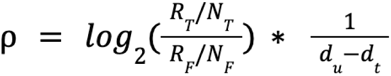

In the GI screen, single sgRNAs with median read counts less than 35 in either orientation (either first or second) in the expression vector in any condition are removed from analysis in all conditions.

Additionally, a pseudocount of 10 is added to each sgRNA’s read count that passes this threshold. Single sgRNA level phenotypes are then determined by averaging the γ or **τ** of each sgRNA paired with all thirteen negative controls in both orientations (26 independent measurements per sgRNA, 52 per gene).

### GO term enrichment

The enrichr function of the gseapy module in python was used to determine GO term enrichment of gene sets from the nominating ATRi screen, GI/eGI maps, and RNAseq datasets. The gene sets used as a reference were “GO_Biological_Process_2021” and “GO_Molecular_Function_2021”. GO terms were considered significantly enriched in a gene set if the -log10 adjusted p-value associated with the term was >=6.

### Calculating genetic interactions

A GI model is generated for each single query sgRNA by quadratic regression of the relationship between all single sgRNA phenotypes and every paired sgRNA phenotype in which the query sgRNA is present (Figure 2A). A genetic interaction for all query paired sgRNA constructs is then calculated based on the deviation of that pair’s phenotype from the model. These differences are then z-score normalized by the standard deviation of the negative control distribution for that guide (standard deviation of phenotypes in the population of 26 query sgRNA-ntc pairs). GIs called for each unique pair of sgRNA are averaged across both orientations of that pair (first and second in the construct). Gene level GIs are called by averaging all four orientations of guides that target those genes.

### Clustering genes by GI profile

GI and eGI matrices were clustered using the “clustermap” function of the seaborn package in Python with the “method” and “metric” variables assigned as “average” and “correlation” respectively. Ontology level clustering as shown in Figure 3A&C is calculated by setting a Pearson distance threshold that maximizes formation of new clusters without collapsing existing clusters together. We qualitatively decided on a Pearson distance of 0.55 as a best fit for all five maps. Similarly, for “System” clusters, we decided on a Pearson distance of 0.8 to sufficiently connect genes to the point where they represented larger areas of biology. To generate the “consensus” clustering, the GI and eGI map were first normalized by their standard deviation, then concatenated to form a 304 by 1520 element matrix. This matrix was clustered using the same methodology as the five independent maps.

### Differential expression analysis of RNAseq data

Differential expression of genes calculated from RNAseq data was analyzed using the DeSeq2 package in R. Genes with fewer than 10 reads across all replicates and conditions were removed from the analysis as they are assumed to not be expressed in either K562 or A549 cells. All comparisons between conditions within a cell line were used to generate differential expression log-fold change and associated p-values.

## References

1. A. Tsherniak, F. Vazquez, P. G. Montgomery, B. A. Weir, G. Kryukov, G. S. Cowley, S. Gill, W. F. Harrington, S. Pantel, J. M. Krill-Burger, R. M. Meyers, L. Ali, A. Goodale, Y. Lee, G. Jiang, J. Hsiao, W. F. J. Gerath, S. Howell, E. Merkel, M. Ghandi, L. A. Garraway, D. E. Root, T. R. Golub, J. S. Boehm, W. C. Hahn, Defining a Cancer Dependency Map. Cell. 170, 564–576.e16 (2017).

2. J. Barretina, G. Caponigro, N. Stransky, K. Venkatesan, A. A. Margolin, S. Kim, C. J. Wilson, J. Lehár, G. V. Kryukov, D. Sonkin, A. Reddy, M. Liu, L. Murray, M. F. Berger, J. E. Monahan, P. Morais, J. Meltzer, A. Korejwa, J. Jané-Valbuena, F. A. Mapa, J. Thibault, E. Bric-Furlong, P. Raman, A. Shipway, I. H. Engels, J. Cheng, G. K. Yu, J. Yu, P. Aspesi, M. de Silva, K. Jagtap, M. D. Jones, L. Wang, C. Hatton, E. Palescandolo, S. Gupta, S. Mahan, C. Sougnez, R. C. Onofrio, T. Liefeld, L. MacConaill, W. Winckler, M. Reich, N. Li, J. P. Mesirov, S. B. Gabriel, G. Getz, K. Ardlie, V. Chan, V. E. Myer, B. L. Weber, J. Porter, M. Warmuth, P. Finan, J. L. Harris, M. Meyerson, T. R. Golub, M. P. Morrissey, W. R. Sellers, R. Schlegel, L. A. Garraway, The Cancer Cell Line Encyclopedia enables predictive modelling of anticancer drug sensitivity. Nature. 483, 603–607 (2012).

3. J. M. Replogle, R. A. Saunders, A. N. Pogson, J. A. Hussmann, A. Lenail, A. Guna, L. Mascibroda, E. J. Wagner, K. Adelman, G. Lithwick-Yanai, N. Iremadze, F. Oberstrass, D. Lipson, J. L. Bonnar, M. Jost, T. M. Norman, J. S. Weissman, Mapping information-rich genotype-phenotype landscapes with genome-scale Perturb-seq. Cell. 185, 2559–2575.e28 (2022).

4. C. Stark, B.-J. Breitkreutz, T. Reguly, L. Boucher, A. Breitkreutz, M. Tyers, BioGRID: a general repository for interaction datasets. Nucleic Acids Res. 34, D535–539 (2006).

5. M. Costanzo, E. Kuzmin, J. van Leeuwen, B. Mair, J. Moffat, C. Boone, B. Andrews, Global Genetic Networks and the Genotype-to-Phenotype Relationship. Cell. 177, 85–100 (2019).

6. C. Boone, H. Bussey, B. J. Andrews, Exploring genetic interactions and networks with yeast. Nat. Rev. Genet. 8, 437–449 (2007).

7. M. Costanzo, B. VanderSluis, E. N. Koch, A. Baryshnikova, C. Pons, G. Tan, W. Wang, M. Usaj, J. Hanchard, S. D. Lee, V. Pelechano, E. B. Styles, M. Billmann, J. van Leeuwen, N. van Dyk, Z.-Y. Lin, E. Kuzmin, J. Nelson, J. S. Piotrowski, T. Srikumar, S. Bahr, Y. Chen, R. Deshpande, C. F. Kurat, S. C. Li, Z. Li, M. M. Usaj, H. Okada, N. Pascoe, B.-J. San Luis, S. Sharifpoor, E. Shuteriqi, S. W. Simpkins, J. Snider, H. G. Suresh, Y. Tan, H. Zhu, N. Malod-Dognin, V. Janjic, N. Przulj, O. G. Troyanskaya, I. Stagljar, T. Xia, Y. Ohya, A.-C. Gingras, B. Raught, M. Boutros, L. M. Steinmetz, C. L. Moore, A. P. Rosebrock, A. A. Caudy, C. L. Myers, B. Andrews, C. Boone, A global genetic interaction network maps a wiring diagram of cellular function. Science. 353, aaf1420 (2016).

8. M. Costanzo, G. Giaever, C. Nislow, B. Andrews, Experimental approaches to identify genetic networks. Curr. Opin. Biotechnol. 17, 472–480 (2006).

9. S. J. Dixon, M. Costanzo, A. Baryshnikova, B. Andrews, C. Boone, Systematic mapping of genetic interaction networks. Annu. Rev. Genet. 43, 601–625 (2009).

10. J. Domingo, P. Baeza-Centurion, B. Lehner, The Causes and Consequences of Genetic Interactions (Epistasis). Annu. Rev. Genomics Hum. Genet. 20, 433–460 (2019).

11. S. R. Collins, K. M. Miller, N. L. Maas, A. Roguev, J. Fillingham, C. S. Chu, M. Schuldiner, M. Gebbia, J. Recht, M. Shales, H. Ding, H. Xu, J. Han, K. Ingvarsdottir, B. Cheng, B. Andrews, C. Boone, S. L. Berger, P. Hieter, Z. Zhang, G. W. Brown, C. J. Ingles, A. Emili, C. D. Allis, D. P. Toczyski, J. S. Weissman, J. F. Greenblatt, N. J. Krogan, Functional dissection of protein complexes involved in yeast chromosome biology using a genetic interaction map. Nature. 446, 806–810 (2007).

12. M. Costanzo, A. Baryshnikova, J. Bellay, Y. Kim, E. D. Spear, C. S. Sevier, H. Ding, J. L. Y. Koh, K. Toufighi, S. Mostafavi, J. Prinz, R. P. St Onge, B. VanderSluis, T. Makhnevych, F. J. Vizeacoumar, S. Alizadeh, S. Bahr, R. L. Brost, Y. Chen, M. Cokol, R. Deshpande, Z. Li, Z.-Y. Lin, W. Liang, M. Marback, J. Paw, B.-J. San Luis, E. Shuteriqi, A. H. Y. Tong, N. van Dyk, I. M. Wallace, J. A. Whitney, M. T. Weirauch, G. Zhong, H. Zhu, W. A. Houry, M. Brudno, S. Ragibizadeh, B. Papp, C. Pál, F. P. Roth, G. Giaever, C. Nislow, O. G. Troyanskaya, H. Bussey, G. D. Bader, A.-C. Gingras, Q. D. Morris, P. M. Kim, C. A. Kaiser, C. L. Myers, B. J. Andrews, C. Boone, The genetic landscape of a cell. Science. 327, 425–431 (2010).

13. B. Fischer, T. Sandmann, T. Horn, M. Billmann, V. Chaudhary, W. Huber, M. Boutros, A map of directional genetic interactions in a metazoan cell. eLife. 4, e05464 (2015).

14. J. A. Doudna, E. Charpentier, Genome editing. The new frontier of genome engineering with CRISPR-Cas9. Science. 346, 1258096 (2014).

15. O. Shalem, N. E. Sanjana, E. Hartenian, X. Shi, D. A. Scott, T. Mikkelson, D. Heckl, B. L. Ebert, D. E. Root, J. G. Doench, F. Zhang, Genome-scale CRISPR-Cas9 knockout screening in human cells. Science. 343, 84–87 (2014).

16. M. Nakamura, Y. Gao, A. A. Dominguez, L. S. Qi, CRISPR technologies for precise epigenome editing. Nat. Cell Biol. 23, 11–22 (2021).

17. A. Pickar-Oliver, C. A. Gersbach, The next generation of CRISPR-Cas technologies and applications. Nat. Rev. Mol. Cell Biol. 20, 490–507 (2019).

18. L. A. Gilbert, M. H. Larson, L. Morsut, Z. Liu, G. A. Brar, S. E. Torres, N. Stern-Ginossar, O. Brandman, E. H. Whitehead, J. A. Doudna, W. A. Lim, J. S. Weissman, L. S. Qi, CRISPR-mediated modular RNA-guided regulation of transcription in eukaryotes. Cell. 154, 442–451 (2013).

19. D. Du, A. Roguev, D. E. Gordon, M. Chen, S.-H. Chen, M. Shales, J. P. Shen, T. Ideker, P. Mali, L. S. Qi, N. J. Krogan, Genetic interaction mapping in mammalian cells using CRISPR interference. Nat. Methods. 14, 577–580 (2017).

20. M. A. Horlbeck, A. Xu, M. Wang, N. K. Bennett, C. Y. Park, D. Bogdanoff, B. Adamson, E. D. Chow, M. Kampmann, T. R. Peterson, K. Nakamura, M. A. Fischbach, J. S. Weissman, L. A. Gilbert, Mapping the Genetic Landscape of Human Cells. Cell. 174, 953–967.e22 (2018).

21. M. Olivieri, T. Cho, A. Álvarez-Quilón, K. Li, M. J. Schellenberg, M. Zimmermann, N. Hustedt, S. E. Rossi, S. Adam, H. Melo, A. M. Heijink, G. Sastre-Moreno, N. Moatti, R. K. Szilard, A. McEwan, A. K. Ling, A. Serrano-Benitez, T. Ubhi, S. Feng, J. Pawling, I. Delgado-Sainz, M. W. Ferguson, J. W. Dennis, G. W. Brown, F. Cortés-Ledesma, R. S. Williams, A. Martin, D. Xu, D. Durocher, A Genetic Map of the Response to DNA Damage in Human Cells. Cell. 182, 481–496.e21 (2020).

22. L. Przybyla, L. A. Gilbert, A new era in functional genomics screens. Nat. Rev. Genet. 23, 89–103 (2022).

23. C. Bock, P. Datlinger, F. Chardon, M. A. Coelho, M. B. Dong, K. A. Lawson, T. Lu, L. Maroc, T. M. Norman, B. Song, G. Stanley, S. Chen, M. Garnett, W. Li, J. Moffat, L. S. Qi, R. S. Shapiro, J. Shendure, J. S. Weissman, X. Zhuang, High-content CRISPR screening. Nat. Rev. Methods Primer. 2, 9 (2022).

24. S. M. Hill, N. K. Nesser, K. Johnson-Camacho, M. Jeffress, A. Johnson, C. Boniface, S. E. F. Spencer, Y. Lu, L. M. Heiser, Y. Lawrence, N. T. Pande, J. E. Korkola, J. W. Gray, G. B. Mills, S. Mukherjee, P. T. Spellman, Context Specificity in Causal Signaling Networks Revealed by Phosphoprotein Profiling. Cell Syst. 4, 73–83.e10 (2017).

25. A. Frost, M. G. Elgort, O. Brandman, C. Ives, S. R. Collins, L. Miller-Vedam, J. Weibezahn, M. Y. Hein, I. Poser, M. Mann, A. A. Hyman, J. S. Weissman, Functional repurposing revealed by comparing S. pombe and S. cerevisiae genetic interactions. Cell. 149, 1339–1352 (2012).

26. M. Costanzo, J. Hou, V. Messier, J. Nelson, M. Rahman, B. VanderSluis, W. Wang, C. Pons, C. Ross, M. Ušaj, B.-J. San Luis, E. Shuteriqi, E. N. Koch, P. Aloy, C. L. Myers, C. Boone, B. Andrews, Environmental robustness of the global yeast genetic interaction network. Science. 372, eabf8424 (2021).

27. S. Bandyopadhyay, M. Mehta, D. Kuo, M.-K. Sung, R. Chuang, E. J. Jaehnig, B. Bodenmiller, K. Licon, W. Copeland, M. Shales, D. Fiedler, J. Dutkowski, A. Guénolé, H. van Attikum, K. M. Shokat, R. D. Kolodner, W.-K. Huh, R. Aebersold, M.-C. Keogh, N. J. Krogan, T. Ideker, Rewiring of genetic networks in response to DNA damage. Science. 330, 1385–1389 (2010).

28. L. A. Gilbert, M. A. Horlbeck, B. Adamson, J. E. Villalta, Y. Chen, E. H. Whitehead, C. Guimaraes, B. Panning, H. L. Ploegh, M. C. Bassik, L. S. Qi, M. Kampmann, J. S. Weissman, Genome-Scale CRISPR-Mediated Control of Gene Repression and Activation. Cell. 159, 647–661 (2014).

29. M. A. Horlbeck, L. A. Gilbert, J. E. Villalta, B. Adamson, R. A. Pak, Y. Chen, A. P. Fields, C. Y. Park, J. E. Corn, M. Kampmann, J. S. Weissman, Compact and highly active next-generation libraries for CRISPR-mediated gene repression and activation. eLife. 5, e19760 (2016).

30. S. Ruiz, C. Mayor-Ruiz, V. Lafarga, M. Murga, M. Vega-Sendino, S. Ortega, O. Fernandez-Capetillo, A Genome-wide CRISPR Screen Identifies CDC25A as a Determinant of Sensitivity to ATR Inhibitors. Mol. Cell. 62, 307–313 (2016).

31. C. Wang, G. Wang, X. Feng, P. Shepherd, J. Zhang, M. Tang, Z. Chen, M. Srivastava, M. E. McLaughlin, N. M. Navone, G. T. Hart, J. Chen, Genome-wide CRISPR screens reveal synthetic lethality of RNASEH2 deficiency and ATR inhibition. Oncogene. 38, 2451–2463 (2019).

32. N. Y. L. Ngoi, M. M. Pham, D. S. P. Tan, T. A. Yap, Targeting the replication stress response through synthetic lethal strategies in cancer medicine. Trends Cancer. 7, 930–957 (2021).

33. H. Goto, T. Natsume, M. T. Kanemaki, A. Kaito, S. Wang, E. C. Gabazza, M. Inagaki, A. Mizoguchi, Chk1-mediated Cdc25A degradation as a critical mechanism for normal cell cycle progression. J. Cell Sci. 132, jcs223123 (2019).

34. T. M. Norman, M. A. Horlbeck, J. M. Replogle, A. Y. Ge, A. Xu, M. Jost, L. A. Gilbert, J. S. Weissman, Exploring genetic interaction manifolds constructed from rich single-cell phenotypes. Science. 365, 786–793 (2019).

35. I. Garcia-Higuera, T. Taniguchi, S. Ganesan, M. S. Meyn, C. Timmers, J. Hejna, M. Grompe, A. D. D’Andrea, Interaction of the Fanconi anemia proteins and BRCA1 in a common pathway. Mol. Cell. 7, 249–262 (2001).

36. K. Townsend, H. Mason, A. N. Blackford, E. S. Miller, J. R. Chapman, G. G. Sedgwick, G. Barone, A. S. Turnell, G. S. Stewart, Mediator of DNA damage checkpoint 1 (MDC1) regulates mitotic progression. J. Biol. Chem. 284, 33939–33948 (2009).

37. A. Ghelli Luserna di Rorà, C. Cerchione, G. Martinelli, G. Simonetti, A WEE1 family business: regulation of mitosis, cancer progression, and therapeutic target. J. Hematol. Oncol.J Hematol Oncol. 13, 126 (2020).

38. F. Merolla, C. Luise, M. T. Muller, R. Pacelli, A. Fusco, A. Celetti, Loss of CCDC6, the first identified RET partner gene, affects pH2AX S139 levels and accelerates mitotic entry upon DNA damage. PloS One. 7, e36177 (2012).

39. Y. Sun, X. Jiang, S. Chen, N. Fernandes, B. D. Price, A role for the Tip60 histone acetyltransferase in the acetylation and activation of ATM. Proc. Natl. Acad. Sci. U. S. A. 102, 13182–13187 (2005).

40. M. Ikura, K. Furuya, S. Matsuda, R. Matsuda, H. Shima, J. Adachi, T. Matsuda, T. Shiraki, T. Ikura, Acetylation of Histone H2AX at Lys 5 by the TIP60 Histone Acetyltransferase Complex Is Essential for the Dynamic Binding of NBS1 to Damaged Chromatin. Mol. Cell. Biol. 35, 4147–4157 (2015).

41. T. Procida, T. Friedrich, A. P. M. Jack, M. Peritore, C. Bönisch, H. C. Eberl, N. Daus, K. Kletenkov, A. Nist, T. Stiewe, T. Borggrefe, M. Mann, M. Bartkuhn, S. B. Hake, JAZF1, A Novel p400/TIP60/NuA4 Complex Member, Regulates H2A.Z Acetylation at Regulatory Regions. Int. J. Mol. Sci. 22, 678 (2021).

42. X. Wang, S. Ahmad, Z. Zhang, J. Côté, G. Cai, Architecture of the Saccharomyces cerevisiae NuA4/TIP60 complex. Nat. Commun. 9, 1147 (2018).

43. D. J. Burgess, J. Doles, L. Zender, W. Xue, B. Ma, W. R. McCombie, G. J. Hannon, S. W. Lowe, M. T. Hemann, Topoisomerase levels determine chemotherapy response in vitro and in vivo. Proc. Natl. Acad. Sci. U. S. A. 105, 9053–9058 (2008).

44. C. Rice, E. Skordalakes, Structure and function of the telomeric CST complex. Comput. Struct. Biotechnol. J. 14, 161–167 (2016).

45. J. Tischler, B. Lehner, A. G. Fraser, Evolutionary plasticity of genetic interaction networks. Nat. Genet. 40, 390–391 (2008).

